# Hydroperoxy-lipids Are Essential Yet Insufficient For Execution Of Ferroptosis

**DOI:** 10.64898/2026.09.11.750967

**Authors:** Sviatlana N. Samovich, Alexander A. Kapralov, Louis J. Sparvero, Brian A. Kleiboeker, Mert Akdogan, Andrew A. Amoscato, Yulia Y. Tyurina, Sally E. Wenzel, Dmitry I. Gabrilovich, Yuri L. Bunimovich, Brent R. Stockwell, Valerian E. Kagan, Hülya Bayır

## Abstract

Cells with irreparable redox disbalance in membranes trigger a program of regulated death – ferroptosis. 15-lipoxygenase (LOX)-catalyzed accumulation of hydroperoxy-phosphatidyl-ethanolamines (HOO-PEs) has been associated with the execution of ferroptosis; yet, experimental proof of their sufficiency is lacking. Using the Fe-independent hydrophobic radical initiator, 2,2’-azobis(2,4-dimethyl)-valeronitrile (AMVN), we demonstrate that the accumulation of hydroperoxy-phospholipids (HOO-PLs) is necessary, but insufficient for driving ferroptosis. We further show that Fe-catalyzed decomposition of HOO-PLs into radical intermediates and oxidatively truncated (OxTr) electrophilic species, as well as the formation of protein adducts with OxTr electrophilic species are required for the completion of the ferroptotic program. Consequently, three types of agents – inhibitors of LOXs and other enzymatic generators of HOO-PLs, radical scavengers, and small-molecule nucleophiles – acting at different stages of lipid peroxidation during the ferroptotic program represent effective and specific ferroptosis regulators and potential therapeutic remedies.

## Introduction

Although life exhibits remarkable adaptability to diverse environmental conditions, the viable range for biological existence remains limited. Extremes of temperature, radiation, pH, oxygen availability, redox potential, and hydrostatic pressure define the physiological boundaries between homeostatic comfort and conditions incompatible with normal cellular functions.^[1,2]^ When stressors exceed the capacity of cells to maintain metabolic integrity and repair damage, intrinsic genetic mechanisms are activated, culminating in regulated death programs (RDPs) that eliminate irreversibly compromised cells.^[3,4]^ Ferroptosis, one of the more recently described RDPs, is initiated when cells fail to maintain redox balance.^[5,6]^ Its execution is associated with an imbalance of thiols and iron, leading to uncontrolled (phospho)lipid peroxidation and resulting in the disintegration of irreparably damaged biomembranes.^[7,8]^

Within just over 13 years since its discovery, ferroptotic cell death has been implicated in the pathogenesis of nearly all major degenerative diseases and acute tissue injuries through mechanisms driven by lipid peroxidation.^[7]^ Formation of hydroperoxy-phospholipids (HOO-PLs), particularly sn-2-hydroperoxy-arachidonoyl-phosphatidylethanolamines (HOO-AA-PEs), is critical for the execution of ferroptotic RDP.^[9,10]^ Conversely, the glutathione (GSH)/ glutathione peroxidase 4 (GPX4)-dependent reduction of HOO-PEs to their corresponding alcohols (HO-PEs) represents a hallmark anti-ferroptotic mechanism that counteracts lipid peroxidation and ferroptotic cell death.^[11,12]^ Similarly, the iPLA_2_β-mediated cleavage of sn-2-hydroperoxyacyl chains from oxidized PE species has been identified as a characteristic anti-ferroptotic mechanism.^[13]^ More recently, phospholipids (PLs) containing oxidizable polyunsaturated fatty acids (PUFA) in both sn-1 and sn-2 positions have been implicated as key drivers of ferroptosis.^[14,15]^ Nevertheless, the capacity of sn-2-HOO-PE species to act as direct executioners of cell death remains experimentally unverified.

Although a potential role for iron (Fe)-catalyzed cleavage of sn-2-hydroperoxy-PEs into radical intermediates, accompanied by their oxidative truncation into electrophilic products, has been proposed,^[16,17]^ this concept remains largely speculative and is based primarily on analogies with hydrocarbon oxidation chemistry rather than on experimentally verified redox mechanisms of ferroptosis. In addition, the role of adducts potentially formed through reactions of oxidatively truncated (OxTr) electrophilic species with nucleophilic sites on proteins remains enigmatic, mainly due to a myriad of possible products and challenges in their identification and quantitative analysis.^[18]^ Clearly, this gap in mechanistic understanding hampers the rational design and development of selective and specific ferroptosis inhibitors or activators with therapeutic potential.

Motivated by these considerations, we utilized LC-MS-based redox phospholipidomics to experimentally delineate the role of key lipid oxidation intermediates in the execution of ferroptosis. We specifically focused on HOO-PLs and corresponding lipid alcohols, free-radical cleavage intermediates, and OxTr electrophilic species that can form adducts with proteins. To experimentally address whether accumulation of HOO-PLs is sufficient to induce ferroptotic death, we employed a strategy to promote phospholipid peroxidation independently of iron supplementation. We employed 2,2′-azobis(2,4-dimethylvaleronitrile) (AMVN), a lipophilic generator of carbon-centered radicals,^[19,20]^ to initiate membrane lipid peroxidation while maintaining basal intracellular iron levels. To assess the contribution of redox-active iron to lipid peroxidation and downstream oxidative fragmentation, we used Fe(III)-nitrilotriacetate (Fe/NTA) as a source of bioavailable iron.^[21]^ We established that HOO-PLs, including HOO-PEs, do not directly cause ferroptotic death. Secondary peroxidation products – free radicals, OxTr species generated from HOO-PLs, and adducts of OxTr electrophilic intermediates with proteins – are essential for the completion of the ferroptotic program. Thus, we propose that three classes of inhibitors – i) inhibitors of LOXs and other enzymatic generators of HOO-PLs, ii) radical scavengers, and iii) small-molecule nucleophiles – may serve as effective regulators of ferroptosis by targeting distinct stages of the lipid peroxidation cascade and preventing the formation of OxTr-protein adducts.

## Results

In ferroptosis, redox-active iron drives the fragmentation of unstable HOO-PLs into reduced (by Fe^2+^) or oxidized (by Fe^3+^) radical intermediates and electrophilic OxTr phospholipid species.^[16,17,22]^ To investigate whether physiological intracellular iron levels are sufficient to support ferroptotic death under conditions of increased HOO-PL formation, HT22 cells were exposed to AMVN, either alone or in combination with Fe/NTA. By assessing cellular susceptibility to death and applying LC-MS-based redox lipidomics to quantify endogenous HOO-PLs and their OxTr cleavage products, we sought to define the role of HOO-PLs in determining a cell’s tolerance and transition to death. In addition, ferroptosis inhibitors, including Ferrostatin-1 (Fer-1) and nucleophilic agents (2-mercaptoethanol (2-ME), dithiothreitol (DTT), and dihydrolipoic acid (DHLA)), were utilized to determine whether blocking radical propagation or trapping OxTr electrophiles affects the susceptibility of HT22 cells to ferroptotic death.

Using redox phospholipidomics, we detected a total of 684 phospholipid species in HT22 cells across all experimental conditions, comprising 388 non-oxidized species (51.5% PUFA-containing species), along with 174 oxygenated lipids and 122 OxTr phospholipids (**Figure 1A, Supplementary Figure S1A**). The distribution of non-oxidized phospholipid subclasses showed that phosphatidylcholine (PC; 58.7 ± 1.8%), was the most abundant, followed by phosphatidylethanolamine (PE; 18.1 ± 1.8%), phosphatidylinositol (PI; 8.4 ± 0.6%), phosphatidylserine (PS; 7.3 ± 0.6%), cardiolipin (CL; 5.9 ± 1.3%), phosphatidic acid (PA; 0.9 ± 0.2%), phosphatidylglycerol (PG; 0.4 ± 0.1%), and bis(monoacylglycero)phosphate (BMP; 0.3 ± 0.1%) (**Figure 1B**). Calculation of the PUFA-to-monounsaturated fatty acid (MUFA)/saturated fatty acid (SFA)-phospholipid ratio revealed a strong predominance of PUFA-containing species, particularly within PI and PE classes (**Supplementary Figure S1B**). Comparison of the total PUFA-containing phospholipids across lipid classes showed the following ranking: PC > PE > PI > PS > PA > BMP > PG (**Figure 1C**). Most of the oxidized species were detected in the PE, PC, PS, PA and PI classes **(Figure 1D**). Oxidation products included species containing one (+1O), two (+2O), three (+3O), or four (+4O) oxygen atoms **(Figure 1D**).

**Figure 1.**
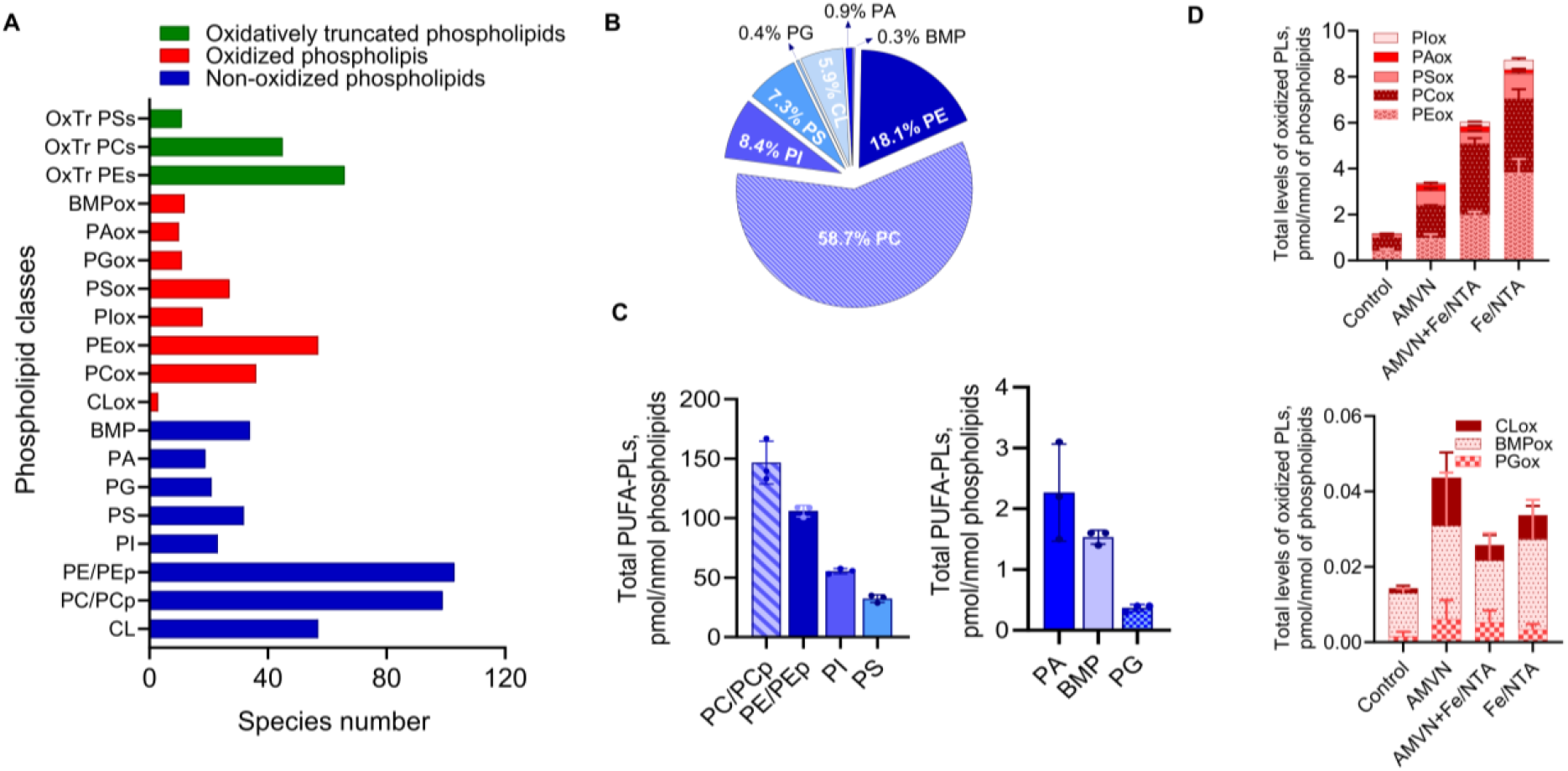
Global LC-MS analysis of the HT22 cell phospholipidome. (A) Number of phospholipid molecular species detected in HT22 cells. 684 phospholipid species, including 388 non-oxidized phospholipids, 174 oxygenated phospholipids and 122 oxidatively truncated phospholipids were identified. PLs, phospholipids CL; cardiolipin, PC, phosphatidylcholine; PE, phosphatidylethanolamine; PS, phosphatidylserine; PI, phosphatidylinositol; PA, phosphatidic acid; PG, phosphatidylglycerol; BMP, bis-monoacylglycero-phosphate; OxTr PLs, oxidatively truncated phospholipids. (B) Pie chart showing the percent distribution of non-oxidized phospholipid subclasses. (C) Bar graphs comparing the total PUFA-containing phospholipids across lipid classes. Data are mean±SD, n = 3/group. Note: two plots are shown to accommodate different abundance ranges. (D) Bar graphs showing total normalized levels of oxidized phospholipids, containing one (+1O), two (+2O), three (+3O), or four (+4O) additional oxygen atoms, across control, AMVN, Fe/NTA, and AMVN+Fe/NTA conditions. Bars are subdivided by lipid class, illustrating the contribution of each class to the overall oxidation. MS^2^ analysis indicates that +1O and +2O species correspond predominantly to hydroxyl (-OH) and hydroperoxy (-OOH) PLs, respectively, which together represent the majority of oxidized species (91.9 ± 3.4 % of total). Data are mean±SD, n = 3-4/group. Note: two plots are shown to accommodate different abundance ranges.

The relative contents of non-truncated and truncated oxidation products are based on semi-quantitative estimates derived from class-based normalization, as described in Methods.

### AMVN Triggers Non-Selective Phospholipid Peroxidation

Because AMVN is a lipophilic compound that incorporates into membrane lipids and spontaneously decomposes to carbon-centered radicals, it is expected to initiate non-enzymatic phospholipid peroxidation proportional to its concentration as well as to the availability of PUFA substrates.^[19,20]^ Based on the known rate constant of AMVN decomposition,^[19]^ we estimated the extent of radical generation and downstream lipid oxidation and found that ~10% of AMVN-derived radicals were associated with the formation of +2O phospholipid species, which predominantly represent HOO-PLs. These values reflect the net accumulation of HOO-PLs under conditions where competing radical-consuming reactions, enzymatic reduction, and downstream conversions also occur.

AMVN treatment alone markedly increased the abundance of oxidized phospholipid species (containing 1-4 oxygens) (**Figure 2A, Supplementary Figure S2A**). Notably, redox lipidomics analysis demonstrated that AMVN induced a 3.6 ± 0.4-fold increase in HOO-PLs compared to control (**Supplementary Figure S2B**). Among all detected HOO-PLs, HOO-PEs (32.3 ± 9.1%), HOO-PCs (29.3 ± 6.0%), and HOO-PAs (29.4 ± 5.4%) together accounted for 91.1% of the total HOO-PLs (**Figure 2, B and C, Supplementary Figure S2C**). As expected, the AMVN induced generation of HOO-PLs was abolished by the radical scavenger Fer-1 (**Figure 2, B and C, Supplementary Figure S2, A-C**).

**Figure 2.**
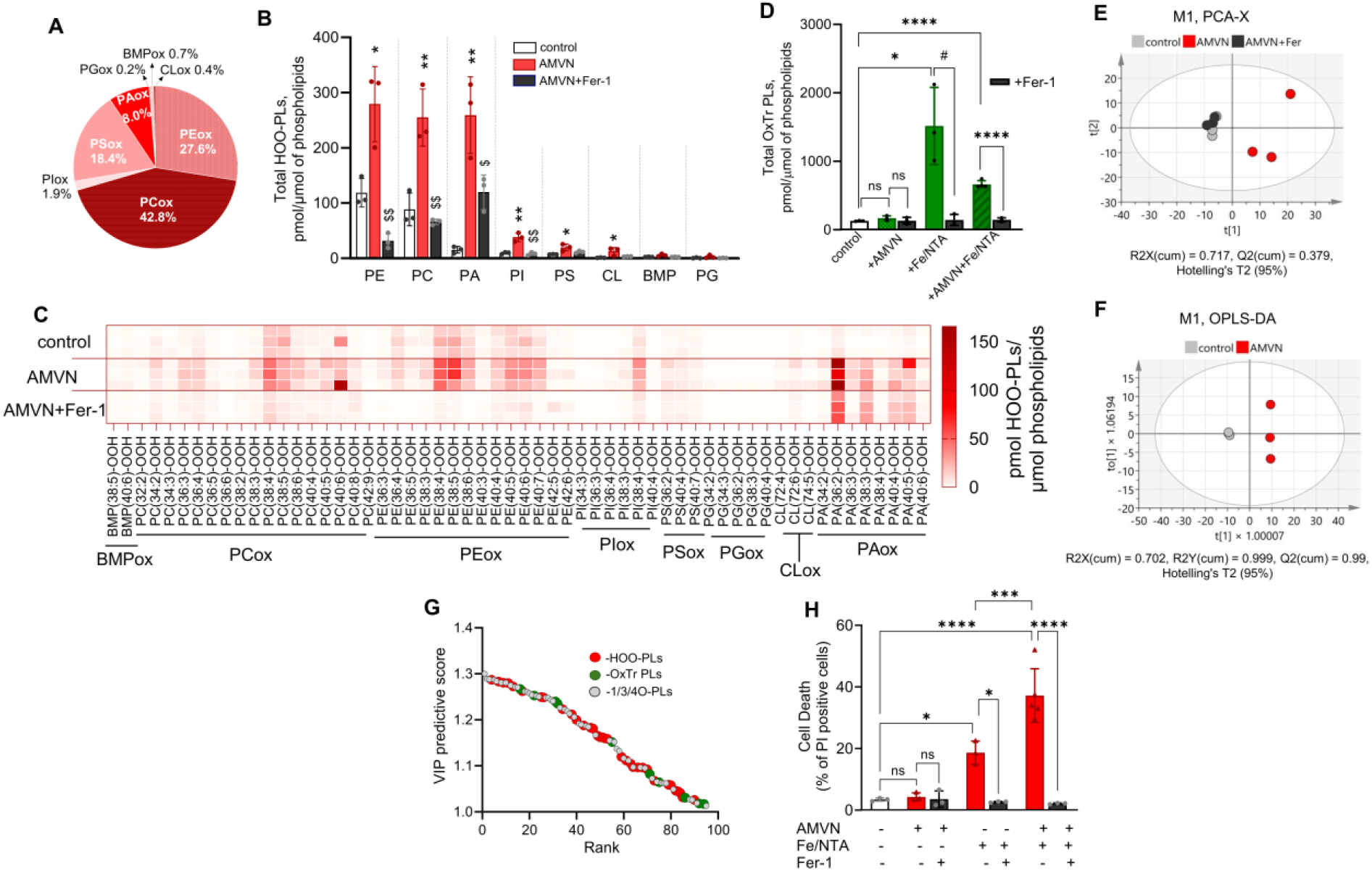
AMVN promotes accumulation of HOO-PLs in HT22 cells independent of ferroptotic death. (A) Pie chart showing the distribution of oxidized phospholipid species (containing one (+1O), two (+2O), three (+3O), or four (+4O) oxygens) across different phospholipid classes upon AMVN treatment. +1O and +2O species, corresponding to OH and OOH modifications by MS^2^, comprise most oxidized lipids (95.3 ± 5.6 %).(B) Quantification of total HOO-PLs (pmol/µmol of phospholipids) across major phospholipid classes under control, AMVN, or AMVN+Fer-1 treatments. Data are mean±SD, n = 3/group. Student t-test, *p < 0.05, **p < 0.01 vs. control; ^$^p < 0.05, ^$$^p < 0.005 vs. AMVN. (C) Heat map of HOO-PL species for control, AMVN, and AMVN+Fer-1 conditions. Data are expressed as pmol/μmol of phospholipids and presented as a heat map (n = 3/group). (D) Total OxTr phospholipid pool (sum of truncated PE, PC, and PS species) measured under control, AMVN, Fe/NTA, and AMVN+Fe/NTA conditions. Data are mean±SD, n = 3-4/group, Student t-test, *p < 0.05, ^#^p < 0.05, ****p < 0.0001. (E) PCA score plot of oxygenated and OxTr phospholipids showing distinct clustering of AMVN-treated cells vs. control, with AMVN+Fer-1 overlapping with control. (F) OPLS-DA score plot confirming clear separation between AMVN and control groups. (G) OPLS-DA-derived VIP predictive score plot (VIPpred > 1) showing the 94 group-separating species in the AMVN treatment. (H) Quantification of cell death under all conditions. Data are mean±SD, n = 3-4/group, Ordinary one-way ANOVA test, *p < 0.05, ***p < 0.001, ****p < 0.0001.

In cells, native pools of redox-active iron can catalyze the cleavage of hydroperoxy-lipids to yield free radicals and OxTr species.^[16,17]^ Because of the short lifetimes of radical intermediates, their contents could not be determined by redox lipidomics. Therefore, we next assessed OxTr phospholipid species in HT22 cells. A total of 122 OxTr species were detected in major phospholipid classes (PE, PC, and PS), with no significant difference in overall OxTr levels between AMVN-treated and control cells (**Figure 2D**).

Accordingly, Fer-1 treatment did not affect OxTr levels in AMVN-treated cells (**Figure 2D**). Only 5 of the 122 individual OxTr species were significantly elevated following AMVN exposure (**Supplementary Figure S2D**), representing a very small fraction of the entire OxTr pool (**Supplementary Figure S3, A-C**). Non-truncated HOO-/HO-PLs represented 98.2 ± 1.7% of the oxidized and truncated phospholipid pool, based on AMVN-specific increases after subtracting control levels.

To further explore the lipidome-wide response, we applied multivariate analysis. Principal component analysis (PCA) of identified oxidized phospholipids and OxTr electrophiles showed that control and AMVN-treated cells formed two well-separated clusters, whereas control and AMVN+Fer-1-treated cells overlapped (**Figure 2E**). Orthogonal projections to latent structures-discriminant analysis (OPLS-DA) confirmed that AMVN treatment strongly affected phospholipid profiles, with tight clustering of controls and clear separation from AMVN-treated cells (**Figure 2F**). The model was validated by CV-ANOVA (p = 0.0152) and permutation testing (200 permutations). To identify the phospholipid species responsible for group separation, we used OPLS-DA–derived S-plot (**Supplementary Figure S4A**) and VIP predictive score plot (**Figure 2G**). The S-plot showed that the hydroperoxy-species, including PC(36:3)+2O, PE(38:4)+2O, and PA (36:3)+2O, were significantly associated with AMVN treatment (**Supplementary Figure S4A**). VIPpred analysis (VIPpred > 1) revealed 94 group-separating species, the majority of which were HOO-PLs (37.2%), while OxTr lipids contributed minimally (10.6%) (**Supplementary Figure S4B**).

Notably, the same three HOO-PLs identified in the S-plot were among the top ten species contributing most to the VIP scores, with no OxTr species appearing in this group (**Supplementary Figure S4C**). Among HOO-PLs with VIPpred > 1, the majority were in PE and PC classes. Overall, these results indicate that HOO-PL species are enriched in AMVN-treated samples and contribute to the observed lipidomic changes.

Importantly, despite extensive accumulation of phospholipid hydroperoxides, AMVN did not induce ferroptotic cell death, regardless of whether Fer-1 was present or absent (**Figure 2H**). Because AMVN treatment promotes predominantly the accumulation of non-truncated lipid peroxidation products, including HO-/HOO-PLs, these data indicate that hydroperoxy-phospholipids alone are not sufficient to trigger ferroptotic cell death.

### Combined Effect Of AMVN/Iron On Phospholipid Peroxidation And Ferroptosis

The initiation of ferroptotic cell death is inextricably linked to iron.^[23]^ The hydroperoxy group in oxidized phospholipids readily undergoes cleavage in the presence of transition metals such as Fe.^[16,17]^ To examine the role of iron in catalyzing the decomposition of hydroperoxy-phospholipids and promoting ferroptosis, HT22 cells were treated with AMVN in combination with Fe/NTA, a bioavailable iron complex that mimics physiologically accessible Fe pools without causing non-specific precipitation.^[21]^ This treatment induced a robust increase in overall phospholipid peroxidation, with a pronounced accumulation of OxTr phospholipids (**Figures 2D and 3A, Supplementary Figure S3, A-C**). The total pool of OxTr species was significantly higher in cells exposed to AMVN+Fe/NTA compared to control cells (5.2 ± 0.6-fold increase) (**Figure 2D**), and these truncated products were detected across major classes of phospholipids (**Figure 3A**). A marked increase in total HOO-PL content (7.3 ± 0.7-fold increase) was also observed (**Figure 3B, Supplementary Figure S5, A-B**), consistent with free radical propagation of lipid peroxidation in the presence of Fe/NTA. It should be noted that Fe^2+^/Fe^3+^-catalyzed lipid peroxidation occurs many orders of magnitude faster than lipid radical generation from AMVN decomposition. The primary products of Fe/NTA reactions are HOO-/HO-PLs, formed via Fe-catalyzed lipid peroxidation of PUFA-PLs,^[16,17,22]^ with HOO-PLs capable of yielding OxTr species.^[16]^ To compare the ferroptotic potential of HOO-PLs under these two conditions, we chose the Fe/NTA concentration to produce levels of HOO/HO-PLs comparable to those generated by AMVN. Indeed, the accumulated levels of HOO/HO-PLs under these two conditions were of the same order of magnitude. In spite of that, the combination of AMVN and Fe/NTA sharply increased cell death, with 37.2 ± 8.7% of HT22 cells affected (an increase of 32.9 ± 10.0% compared to AMVN alone) (**Figure 2H**). Notably, this cell death was completely blocked by Fer-1 (**Figure 2H**), which also substantially prevented the accumulation of OxTr PLs (**Figures 2D and 3A, Supplementary Figure S3, A-C**) and reduced HOO-PL levels (**Figure 3B, Supplementary Figure S5, A-B**). These results demonstrate that accumulation of non-truncated HOO/HO-PL species under AMVN alone is not sufficient to trigger ferroptosis, whereas the addition of Fe/NTA promotes secondary lipid peroxidation, including the formation of OxTr species, and strongly enhances ferroptotic cell death.

**Figure 3.**
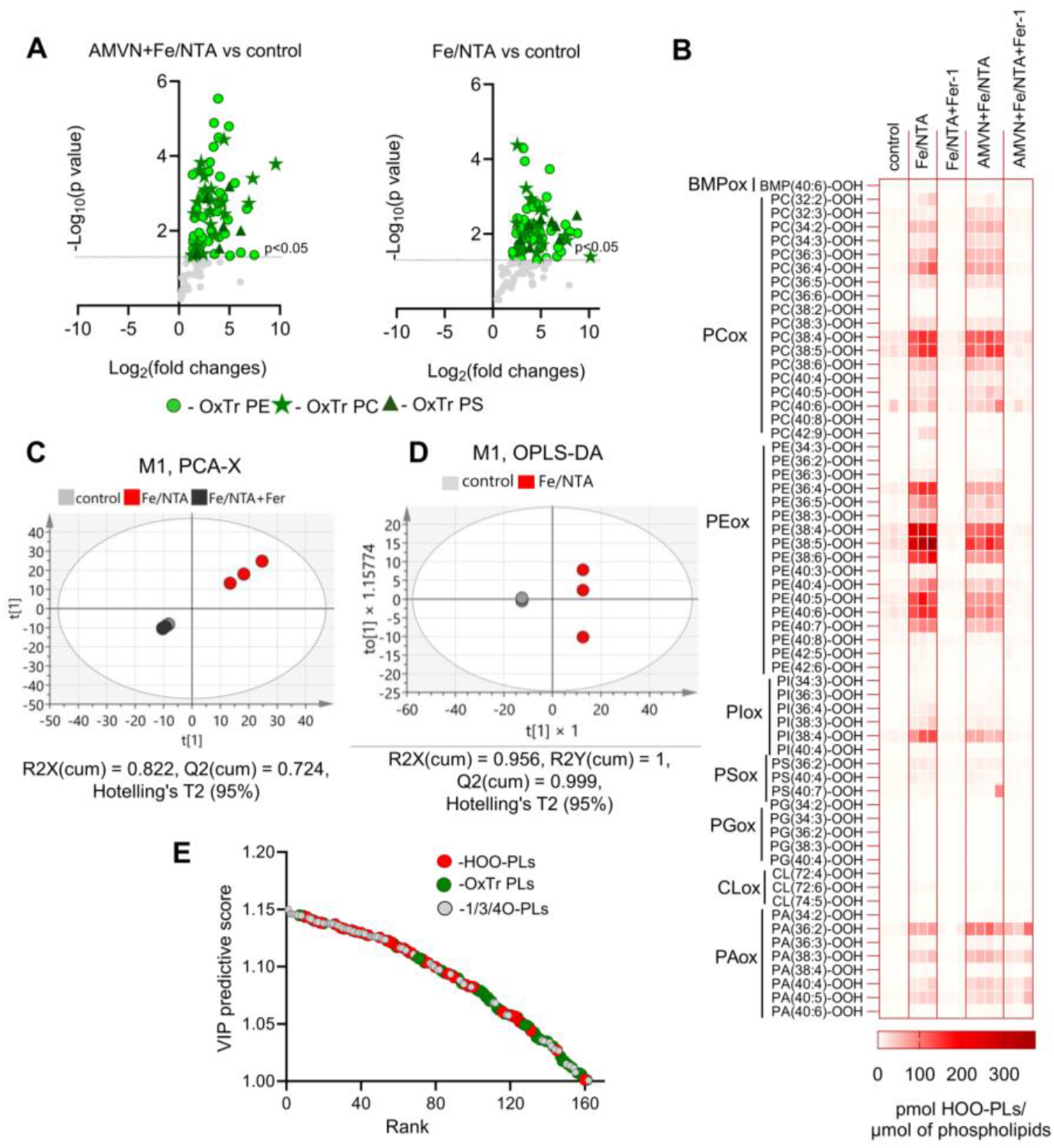
Oxidized and oxidatively truncated phospholipid profiles and multivariate analyses of AMVN+Fe/NTA- and Fe/NTA-treated HT22 cells. (A) Volcano plots showing significantly altered OxTr phospholipid species (truncated PE, PC, and PS) for (left) AMVN+Fe/NTA-treated vs. control cells and for (right) Fe/NTA-treated vs. control cells. (B) Heat map of HOO-PL species for control, Fe/NTA, Fe/NTA +Fer-1, AMVN+Fe/NTA and AMVN+Fe/NTA +Fer-1 conditions. Data are expressed as pmol/μmol of phospholipids and presented as a heat map (n = 3/group). (C) PCA score plot of oxidized and OxTr phospholipids showing distinct clustering of Fe/NTA-treated cells vs. control, with Fe/NTA+Fer-1-treated cells overlapping with control. (D) OPLS-DA score plot confirming clear separation between Fe/NTA and control groups within the 95% confidence ellipse. (E) OPLS-DA-derived VIP predictive score plot (VIPpred > 1) showing the 162 group-separating species in the Fe/NTA treatment.

Testing the effects of Fe/NTA alone on HT22 cells revealed a significant increase in oxidized phospholipid species (**Figures 2D, 3A, and 3B, Supplementary Figure S5, B-C**), including a 10.7 ± 1.6-fold accumulation of HOO-PLs relative to control, along with substantial production of OxTr phospholipid products (11.9 ± 0.9-fold increase). These peroxidation effects of Fe/NTA were effectively suppressed by Fer-1 (**Figures 2D, 3A, and 3B, Supplementary Figure S5, B-C**). Fe/NTA treatment alone caused a moderate increase in cell death (15.2 ± 4.3% compared to control) (**Figure 2H**), which was significantly smaller than that observed with AMVN+Fe/NTA. Notably, this cell death was completely quenchable by Fer-1 **(Figure 2H**), indicating its ferroptotic nature. To uncover oxidized phospholipid metabolites driving ferroptosis in Fe/NTA-treated HT22 cells, we applied multivariate PCA (**Figure 3C**) and OPLS-DA (**Figure 3D**). S-plot (**Supplementary Figure S5D**) and VIPpred analyses (**Figure 3E**) identified OxTr species, including OxTr-PC(25:0)+2O and OxTr-PE(25:1)+2O, as among the top contributors to group separation (**Supplementary Figure S5, E-F**). Notably, OxTr phospholipids were more prevalent than HOO-PLs (36.4% versus 24.7%, respectively) (**Supplementary Figure S5E**) and emerged as major contributors to the group difference, suggesting that Fe/NTA treatment preferentially promotes the formation of OxTr species over HOO-PL species.

In contrast to AMVN alone, both AMVN+Fe/NTA and Fe/NTA treatments significantly increased the OxTr phospholipid pool compared to control (**Figure 2D**), and this increase was markedly attenuated by Fer-1. To get a better insight into the role of individual OxTr phospholipids as ferroptosis inducers, we performed species-specific univariate analysis, which showed that AMVN+Fe/NTA and Fe/NTA treatments caused significant increases in 81 and 72 OxTr species, respectively, out of the 122 detected. Comparison of significantly altered species across AMVN, Fe/NTA, and AMVN+Fe/NTA treatment groups revealed both unique and shared truncated lipids (**Supplementary Figure S5G**). Interestingly, among OxTr phospholipids common to AMVN+Fe/NTA and Fe/NTA treatments, the majority (55.1%) were truncated PE species, whereas truncated PC and PS species accounted for 30.6% and 14.3%, respectively. Formation of truncated species was confirmed by MS/MS analysis of PE, PC, and PS species, with representative spectra shown in **Supplementary Figure S4, A-C**, exemplifying the characteristic fragmentation patterns and diagnostic ions used to identify these lipids (**Supplementary Figure S6, A-C**). Detailed annotations are summarized in **Supplementary Tables S1–S3**. The ratio of HOO-PLs to OxTr species was high with AMVN alone (5.2 ± 0.8) and decreased markedly in the presence of AMVN+Fe/NTA and Fe/NTA (2.7 ± 0.1 and 1.7 ± 0.4, respectively), highlighting the shift from (hydro)peroxidation to oxidative lipid truncation driven by Fe-dependent mechanisms.

Next, we calculated correlations between total OxTr phospholipid levels and cell death under three conditions (**Supplementary Figure S7A**). No significant correlation was observed in AMVN-treated ± Fer-1 HT22 cells (Pearson r = −0.48, p = 0.18). In contrast, AMVN+Fe/NTA- and Fe/NTA-treated ± Fer-1 HT22 cells exhibited a strong positive correlation between OxTr phospholipids and cell death (AMVN+Fe/NTA: Pearson r = 0.94, p < 0.0001; Fe/NTA: Pearson r = 0.84, p = 0.0043).

To further examine the link between HOO-PLs and ferroptosis, oxidized phospholipid species containing hydroperoxy groups were normalized to ferroptosis-attributable cell death events, defined as the fraction of cell loss rescued by Fer-1. For each condition, Fer-1-sensitive HOO-PL signal was calculated by subtracting levels measured in the presence of Fer-1 from those measured in its absence. Each resulting HOO-PL value was then divided by the number of Fer-1-rescued cells to obtain a normalized, semi-quantitative index of “HOO-PL burden per ferroptotically dying cell.” This normalization revealed condition-dependent differences: AMVN treatment caused the highest HOO-PL accumulation, Fe/NTA produced less, and AMVN+Fe/NTA combination showed the lowest signal (**Supplementary Figure S7B**). Notably, cell death followed the opposite trend, with the AMVN+Fe/NTA condition causing greatest loss of viability (**Figure 2H**).

These results indicate that HOO-PL accumulation alone is insufficient to trigger cell death. Instead, HOO-PLs act as precursors that undergo degradation into oxy-radicals, truncated electrophilic phospholipids, and their protein adducts. These secondary species likely drive the execution of ferroptotic cell death. The absence of ferroptosis in AMVN-treated cells indicates that intracellular mechanisms driven by endogenous iron pools were inactive under these conditions.

### Oxidatively Truncated Phospholipid–Mediated Protein Modification Drives Ferroptosis

The increase in the percentage of ferroptotic cells in the AMVN+Fe/NTA treatment was associated with elevated levels of electrophilic OxTr species (containing oxo- and carboxy-groups) capable of forming covalent adducts with proteins. Consistent with this, truncated PE, PC, and PS species were identified in the AMVN+Fe/NTA group, although their total abundance was 2.3 ± 0.6-fold lower than in the Fe/NTA group (**Figure 2D**). This suggests that formation of adducts between OxTr phospholipids and proteins may contribute to enhanced ferroptotic cell death. Indeed, cell death in AMVN+Fe/NTA-treated cells was 18.6 ± 4.5% greater than in Fe/NTA-treated cells (**Figure 2H**), supporting the idea that protein-OxTr electrophile adducts contribute to ferroptosis execution.

Cleavage of HOO-PLs can result in oxidative modification of proteins by OxTr products through the formation of adducts and cross-linked protein oligomers. Recently, we described such “oxidative phospholipidation”, including the formation of PE-protein adducts during ferroptosis,^[18]^ which can be prevented by radical scavengers, e.g., Fer-1. This raises the question: which of the two types of reactions – radical-dependent cleavage of HOO-PLs or electrophilic addition of OxTr phospholipid species – is required for the execution of ferroptotic death. To address this question, we assessed ferroptotic death and redox lipidomics of HT22 cells treated with AMVN+Fe/NTA in the presence of exogenous low-molecular-weight thiols (2-ME, DTT, and DHLA) (**Supplementary Figure S8A**). We found that all three compounds reduced ferroptotic cell death, with 2-ME displaying the strongest effect (0.5 or 3 mM reduced cell death from 51.9 ± 7.1% to 29.4 ± 4.9% and 14.9 ± 2.8%, respectively) (**Figure 4A**). Notably, these protective effects were comparable to those observed with Fer-1 (17.2 ± 2.8%) **(Figure 4A**). DTT and DHLA also decreased AMVN+Fe/NTA-induced cell death to 25.1 ± 1.4% and 26.8 ± 4.2%, respectively, compared to 40.8 ± 6.0% in the AMVN+Fe/NTA control group (**Figure 4B**). To get an insight into oxidative protein modifications, cell lysates were analyzed by reducing SDS-PAGE followed by silver staining (**Supplementary Figure S8B**). AMVN+Fe/NTA treatment resulted in the appearance of diffuse staining in the high-molecular-weight region (>250 kDa), consistent with the formation of detergent-resistant protein oligomers (**Figure 4A, Supplementary Figure S8B**), likely reflecting protein aggregation driven by intermediates of phospholipid peroxidation. In the presence of 2-ME, cells treated with AMVN+Fe/NTA displayed markedly reduced levels of this oligomerization (**Figure 4A, Supplementary Figure S8B**). Consistent with this, redox lipidomics showed that DTT and DHLA decreased total OxTr PE levels by 51.3% and 44.2%, respectively (**Figure 4C**). Thus, small nucleophiles may “outcompete” protein nucleophilic sites in reactions with OxTr electrophilic peroxidation products, thereby limiting protein modification. This interpretation is consistent with the fact that reactions of lipid oxy-radicals with thiols yield highly reactive pro-oxidant thiyl radicals capable of propagating lipid peroxidation.^[24,25]^ This indicates that strong exogenous small-molecule nucleophiles can protect proteins from oxidative aggregation, thus preventing their pro-ferroptotic lipidation.^[24,26]^

**Figure 4.**
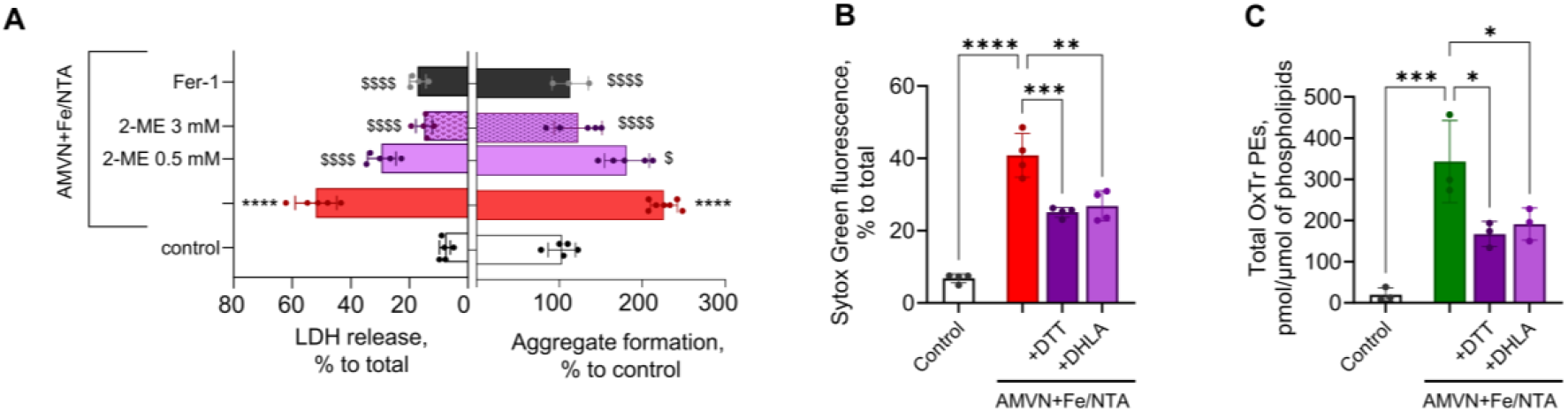
Nucleophile-mediated protection against truncated lipid-protein adducts in HT22 cells. (A) (*Left*) Quantification of cell death measured by LDH release (% of total) in control, AMVN+Fe/NTA, AMVN+Fe/NTA + 2ME (0.5mM), AMVN+Fe/NTA + 2ME (3mM), and AMVN+Fe/NTA + Fer-1 conditions. Data are mean±SD. (*Right*) Quantification of protein aggregate formation (% of total) in control, AMVN+Fe/NTA, AMVN+Fe/NTA + 2ME (0.5mM), AMVN+Fe/NTA + 2ME (3mM), and AMVN+Fe/NTA + Fer-1 conditions. Data are mean±SD, n = 3-7/group, Ordinary one-way ANOVA test, ****p < 0.0001 vs. control; ^$^p = 0.0195, ^$$$$^p < 0.0001 vs. AMVN+Fe/NTA. (B) Quantification of cell death measured by Sytox Green fluorescence (% of total) in control, AMVN+Fe/NTA, AMVN+Fe/NTA + DTT, and AMVN+Fe/NTA + DHLA conditions. Data are mean±SD, n = 4/group, Ordinary one-way ANOVA test, **p < 0.005, ***p < 0.001, ****p < 0.0001. (C) Levels of total OxTr PEs in control, AMVN+Fe/NTA, AMVN+Fe/NTA + DTT, and AMVN+Fe/NTA + DHLA conditions. Data are mean±SD, n = 3/group, Ordinary one-way ANOVA test, *p < 0.05, ***p < 0.005.

## Discussion

“Ferroptosis is a non-apoptotic mechanism of cell death that emerges from the interaction between iron, oxygen, and oxidizable phospholipids (PLs). Compared to other forms of cell death, ferroptosis is uniquely distinguished by the accumulation of *(phospho)lipid hydroperoxides*”.^[7]^ This quote from a recent comprehensive review reflects a widely shared view that elevated lipid hydroperoxide levels are a hallmark of ferroptotic cell death. In contrast, we demonstrate here that although HOO-PLs are necessary for ferroptosis, they are not sufficient for the execution of ferroptotic death.

With the emergence of aerobic life, Fe^2+^ ions in aqueous environments were oxidized to Fe^3+^ and largely precipitated, limiting radical-generating reactions.^[27]^ Yet, organisms require metabolically sufficient iron, which is delivered via sophisticated regulatory mechanisms, including storage proteins and chaperones,^[28-31]^ to prevent unwanted redox activity that could cleave HOO-PLs into radicals and generate electrophilic OxTr species. Dysregulation of these iron-regulatory systems can lead to uncontrolled phospholipid peroxidation, formation of OxTr species, and ultimately ferroptotic cell death.^[16,17,22]^ Clinical interventions, such as intravenous iron therapy, further illustrate how perturbations in iron homeostasis can increase catalytic iron levels^[32-34]^ and predispose cells to ferroptosis.^[35,36]^ Similarly, in cancer and other pathophysiological contexts, iron availability is modulated by proteins such as lipocalin 2 and its receptor, which can influence susceptibility to ferroptotic cell death.^[37-39]]^

Free radical reactions are characterized by the generation of numerous reactive intermediates from unstable –O–O– bonds, particularly in the presence of transition metals like Fe and Cu.^[16,17,22]^ Failure of iron-regulatory mechanisms leads to excessive formation of reactive intermediates and dysregulated phospholipid peroxidation in cellular membranes, yielding a heterogeneous spectrum of OxTr products.^[16,22,40]^ For example, oxidative decomposition of peroxidized arachidonic acid yields >300 products bearing oxygenated electrophilic groups that are either retained or released as leaving groups.^[41,42]^ Given the ability of these intermediates to react with and modify nucleophilic targets in proteins and other macromolecules, one can imagine that a comprehensive analysis of products generated during free radical oxidation of lipids represents an unsurmountable task. A typical example of such metabolic redox dysregulation is ferroptosis.^[7]^ Recent advances in high-resolution mass spectrometry have enabled the detection of ferroptosis-related oxidatively modified lipids and phospholipids in biological systems.^[18,43-45]^ A cautionary note is warranted for non-specific methodologies commonly employed to characterize ferroptosis. One such protocol relies on the oxidation of non-lipid fluorogenic molecules such as BODIPY C11.^[46,47]^ While technically simple, this method – erroneously and misleadingly referred to as “lipid ROS” – can, at best, be utilized merely for correlative studies of pro-oxidant redox environments.^[46]^ However, it does not provide information on the chemistry or biochemistry of lipids and their oxidation products.^[7,47]^ In contrast, LC-MS–based protocols of redox lipidomics enables detailed characterization of a variety of primary and secondary molecular products of lipid peroxidation.

However, their accurate quantification remains challenging due to the limited availability of appropriate deuterated standards, which constrains broader application.^[48,49]^ In practice, measurements are often semi-quantitative and based on relative signal intensities rather than absolute concentrations.

Redox lipidomics analysis established HOO-PLs as primary products of lipid peroxidation generated during ferroptosis.^[9,16,44]^ Both enzymatic (biochemical) and non-enzymatic (chemical) mechanisms are engaged in lipid alkyl radical (L•) generation via a peroxyl radical (LOO•) abstraction of a bis-allylic hydrogen from a non-oxidized lipid molecule (LH).^[16,40]^ High substrate selectivity of enzymatically-catalyzed reactions is regulated by 3D-organization of their catalytic sites. A typical example is the catalytic involvement of LOXs, which effectively oxidize PUFA-PEs but not PUFA-PCs due to their bulky head groups.^[15,50]^ In contrast, chemical lipid peroxidation is randomly driven, with the reaction rate proportional to the number of bis-allylic hydrogens in the oxidizable substrate.^[40]^ The complex of 15LOX with phosphatidylethanolamine binding protein 1 (PEBP1) defines the selectivity and positional specificity of HOO-PE formation,^[51]^ with sn2-15-HOO-arachidonoyl- and sn2-17-HOO-adrenoyl-PE as predominant products.^[9]^ The biological significance of this enzymatic selectivity and specificity is not completely deciphered but may enable precise regulation of the pathway, either through modulation of the 15LOX/PEBP1 complex or by enzymatic reduction of these HOO-PEs to their hydroxy derivatives.

Enzymatic reduction of HOO-PE is most often carried out by GPX4, utilizing GSH as a source of reducing equivalents.^[11,12]^ Alternative reducing mechanisms may be utilized where reactive vicinal sulfhydryls of thioredoxins^[52]^ or dihydrolipoate^[53,54]^ are employed. Importantly, in the absence of enzymatic control (e.g., by GPX4^[12]^ or GPX1^[55]^), thiol-dependent scavenging of lipid radicals (peroxyl and alkoxyl) cannot be considered a protective anti-ferroptotic mechanism, because the resulting thiyl radicals – characterized by highly positive redox potentials^[56]^ – can propagate, rather than quench, phospholipid peroxidation.^[24]^

Mechanistically, AMVN-induced lipid radicals alone yielded robust HOO-PL accumulation but failed to trigger ferroptosis, underscoring that hydroperoxide accumulation is insufficient for cell death. Despite the presence of endogenous pools of redox-active, “loosely bound” intracellular iron capable of decomposition HOO-PLs into secondary lipid peroxidation products, AMVN-derived HOO-PLs did not undergo breakdown.

Supplementation with Fe/NTA facilitated HOO-PL cleavage into OxTr species, amplified radical propagation, and significantly increased ferroptotic cell death. Interestingly, AMVN+Fe/NTA caused greater cell death despite a lower total OxTr abundance than Fe/NTA alone, suggesting that reactive OxTr species and their covalent adducts with vulnerable target proteins, rather than total OxTr levels, are the critical executioners of ferroptosis. The substantial structural diversity of HOO-PL-derived intermediates highlights the need for further exploration to fully elucidate their roles. LO• and LOO• radicals undergo competing intra- and intermolecular reactions, including β-scission, generating two types of electrophilic products: diffusible small molecules and membrane- or protein-retained truncated lipids. At present, redox lipidomics does not permit accurate discrimination and quantification of these highly diverse intermediates. However, several speculations can be made. Importantly, AMVN contributes to the accumulation of HOO-PLs, thereby increasing the pool of substrates available for Fe-mediated cleavage and generation of electrophilic products. This mechanism may represent a primary driver of the enhanced formation of reactive OxTr species and the increased ferroptotic cell death observed under AMVN+Fe/NTA conditions. Moreover, in the absence of AMVN, Fe/NTA may favor the formation of OxTr phospholipid alcohols from HOO-PLs through Fe^2+^-mediated reductive cleavage, generating alkoxyl radicals (LO•), a process that proceeds more rapidly than Fe^3+^-mediated peroxyl radical (LOO•) formation.^[57]^ In contrast, in the presence of AMVN, its spontaneous decomposition generates carbon-centered radicals that react with molecular oxygen to form peroxyl radicals.^[19]^ These radicals can either yield highly reactive electrophilic OxTr phospholipid aldehydes in an iron-independent manner or propagate lipid peroxidation in an iron-dependent fashion, generating new HOO-PLs and amplifying the overall peroxidation process, thus explaining the more pronounced cell death observed under AMVN+Fe/NTA conditions. Consequently, preferential Fe-catalyzed formation of less toxic OxTr phospholipid alcohols under a reducing environment (Fe/NTA alone), versus generation of reactive OxTr phospholipid aldehydes and protein adduct accumulation (AMVN+Fe/NTA) may account for the lower abundance of OxTr intermediates yet higher levels of ferroptotic cell death observed in the AMVN+Fe/NTA conditions.

The present study demonstrates that potent small-molecule nucleophiles can act as anti-ferroptotic agents by preventing electrophilic modification of proteins by OxTr phospholipid species. In this context, thiols function primarily as two-electron nucleophiles, reacting with electrophilic OxTr species (e.g., via Michael addition), rather than as classical radical-scavenging agents. Incomplete suppression of cell death by nucleophiles (2-ME, DTT, and DHLA) indicates the contribution of additional radical-based pro-ferroptotic mechanisms unrelated to OxTr species.^[24]^ In this context, the well-established protective roles of thioredoxins and other sulfhydryl-rich proteins (e.g., metallothioneins) should be taken into account.^[52,58]^

## Conclusion

Our study demonstrates that HOO-PLs, long considered hallmarks of ferroptosis, are necessary but not sufficient for the execution of the death program. Using redox phospholipidomics and an Fe-independent radical initiator (AMVN), we show that ferroptosis requires a sequential process: initial accumulation of HOO-PLs, iron-dependent generation of reactive radicals and electrophilic OxTr lipids, penultimately leading to protein modification. These findings resolve a key conceptual gap in ferroptosis biology and establish a mechanistic framework that distinguishes lipid hydroperoxide formation from the lethal steps of ferroptotic cell death.

## Supporting Information

Additional experimental details, materials, and methods, including figures and tables as mentioned in the text (PDF).

## Acknowledgements

This work was supported by NIH (CA266342, CA272946, NS076511, HL174611, NS061817).

## Notes

Conflict of Interests Statement: Dmitry I. Gabrilovich is an employee and shareholder of AstraZeneca. The other authors declare no conflict of interest.

### Competing Interest Statement

Dmitry I. Gabrilovich is an employee and shareholder of AstraZeneca. The other authors declare no conflict of interest.

