## Supporting Information for "Hydroperoxy-lipids Are Essential Yet Insufficient For Execution Of Ferroptosis"

**Table of Contents for Supporting Information**


I. Table of Contents 2

II. Experimental Methods 3-7

III. Supplementary Tables S1-S3 7-11

IV. Supplementary Figures S1-S8 11-21

### METHODS

**Materials.** 2,2'-azobis(2,4-dimethyl)-valeronitrile (AMVN, QC-2185) was purchased from Combi-Blocks (San Diego, CA). 2-mercaptoethanol (2-ME, M6250), dihydrolipoic acid (DHLA, 437694), iron(III) chloride (157740), nitrilotriacetic acid (N9877), and 2,6-di-tert-butyl-4-methylphenol (BHT, 128370) were obtained from Sigma-Aldrich. All deuterated phospholipids used as reference standards were purchased from Avanti Polar Lipids (Alabaster, AL, USA). Ferrostatin-1 (Fer-1, S7243) was obtained from Selleck Chemicals. Unless otherwise stated, all other reagents were purchased from Fisher Scientific (Waltham, MA).

**Cell culture.** Mouse Hippocampal Neuronal Cell Line HT22 (Sigma, SCC129) was maintained in Dulbecco's modified Eagle's medium (DMEM, Gibco 11995073) with 10 % FBS, 1X penicillin/streptomycin and incubated at 37 °C, 5% $CO_2$ and 95% humidity.

**Ferroptosis assay.** HT22 cells were seeded in 48-well plates in DMEM medium with 0.18 mM AMVN and/or 0.3 mM Fe/NTA for 18h in the presence or absence of Fer-1 (0.4μM), DTT (3mM), DHLA (3mM), or 2-ME (0.5mM or 3mM). Cell death was assessed by flow cytometry. After incubation, cells were trypsinized, centrifuged at 700 × g for 6 min, resuspended in PBS containing propidium iodide (PI) for 5 min on ice, and then monitored by flow cytometry using FACS Canto flow cytometer (BD Bioscience). Data were analyzed using FlowJo Software (FlowJo. LLC). For lipidomics analysis, HT22 cells were seeded at $1.4 \times 10^6$ cells per dish (15 cm) and incubated in DMEM medium with 0.18 mM AMVN and/or 0.3 mM Fe/NTA for 18h in the presence or absence of Fer-1 (0.4μM), DTT (3mM) or DHLA (3mM).

Cell death was also determined by lactate dehydrogenase (LDH) release and Sytox Green staining. *LDH assay.* LDH activity was quantified using the Cytotoxicity Detection Kit (LDH) according to the manufacturer's instructions (Promega Corporation, Madison, WI). *Sytox Green Staining.* After 24 h of incubation, 2 μM Sytox Green was added to stain dead cells. Fluorescence was measured using a Cytation 5 imaging reader (excitation/emission: 500/530 nm). To quantify total Sytox Green fluorescence, cells were lysed with 0.1% Triton X-100.

**Detection of high-molecular-weight detergent-resistant protein species.** HT22 cells were seeded at $1.4 \times 10^6$ cells per 15 cm dish and treated with 0.18 mM AMVN and/or 0.3 mM Fe/NTA for 18 h. Cells were harvested, washed with PBS, and lysed in RIPA buffer supplemented with protease inhibitors. Protein concentrations were determined using Thermo Scientific Pierce BCA Protein Assay Kit, and equal amounts of protein (e.g., 25 μg per lane) were resolved by SDS-PAGE on 4-16% gradient gels. Proteins were visualized using Thermo Scientific Pierce Silver

Stain Kit according to the manufacturer's instructions. High-molecular-weight, detergent-resistant material appearing as unresolved signal above 250 kDa was quantified by densitometric analysis using Image J software. Signal intensity within this region of interest was normalized to the corresponding region in control samples to account for loading variability. For the purpose of this study, unresolved high-molecular-weight material was operationally defined as aggregated protein species. Data represent 3-7 independent biological replicates.

**Lipid extraction.** Lipids were extracted from HT22 cells using the Folch procedure[59] and phosphorus[60] was determined by a micro-method. Briefly, cells (1.2-1.5x$10^6$) were resuspended in 0.75% KCl and lipids were extracted using a chloroform methanol mixture of 2:1 (v/v). To prevent oxidation of lipids during extraction and sample preparation for LC-MS analysis, a chloroform-methanol mixture containing 0.01% butylated hydroxytoluene (BHT) was used.

**Phospholipidomics and redox phospholipidomics LC-MS/MS analysis in HT22 cells.** To identify and quantitatively analyze non-oxidized, oxidized, and oxidatively truncated phospholipid species, we performed targeted redox phospholipidomics using LC-ESI/MS. Class-specific separation and high-resolution detection of both non-oxidized and oxidized phospholipids was performed using a Thermo HPLC system coupled to an Orbitrap Fusion Lumos mass spectrometer (Thermo Fisher Scientific). Given the complexity and diversity of oxidatively modified phospholipids, normal-phase chromatography was applied to separate phospholipids by polar head group class, thereby reducing the complexity of co-eluting species and enhancing confidence in class-specific identification. Separations were carried out on a silica column (Luna 3 μm Silica (2), 100 Å, 150 × 1.0 mm; Phenomenex) at a flow rate of 0.065 mL/min. The column was maintained at 35 °C. The analysis was performed using gradient solvents A and B containing 10 mM ammonium formate, as previously described.[13] Solvent A contained isopropanol/hexane/water (285:215:5, v/v/v), and solvent B contained isopropanol/hexane/water (285:215:40, v/v/v). All solvents were LC/MS grade. The gradient was as follows:0–3 min, 10–37% B; 3–15 min, hold at 37% B; 15–23 min, 37–100% B; 23–75 min, hold at 100% B; 75–76 min, 100-10% B; 76–90 min, equilibrate at 10% B. Lipids were analyzed in negative ion mode using the following parameters: capillary voltage, 3500 V; sheath, aux and sweep gases (35, 17, 0, respectively); ion transfer tube temperature, 300 °C; Orbitrap resolution, 120,000; scan range m/z 400-1800. Data-dependent MS/MS was performed using an isolation window of 1.2 m/z and higher-energy collisional dissociation (HCD) with a normalized collision energy of 24. MS/MS spectra were acquired at an Orbitrap resolution of 15,000.

**Lipid identification and in-house database annotation.** Data processing was performed using Compound Discoverer™ 2.0 software (Thermo Fisher Scientific, San Jose, CA) with an in-house-developed workflow and a curated database of non-oxidized, oxidized, and oxidatively truncated phospholipids. The database contains exact masses for common phospholipid species and was internally validated using authentic standards and characteristic MS/MS spectral features, consistent with established redox phospholipidomics workflows.[48] Peaks with a signal-to-noise ratio of > 3 were extracted and searched against the phospholipid database. Lipid species were assigned based on three criteria: retention time (relative to class-specific internal standards), accurate mass (m/z), and MS/MS fragmentation patterns consistent with expected structural features. Mass tolerance was set to 7 ppm with a minimum intensity threshold of 5000. Class-specific retention times were determined using exogenously added deuterated internal standards: 1-hexadecanoyl(d31)-2-(9Z-octadecenoyl)-sn-glycero-3-phospho-ethanolamine (PE(16:0D31/18:1)), 1-hexadecanoyl(d31)-2-(9Z-octadecenoyl)-sn-glycero-3-phosphocholine (PC(16:0D31/18:1)), 1-hexadecanoyl(d31)-2-(9Z-octadecenoyl)-sn-glycero-3-phosphoserine (PS(16:0D31/18:1)), 1-hexadecanoyl(d31)-2-(9Z-octadecenoyl)-sn-glycero-3-phosphate (PA(16:0D31/18:1)), 1-hexadecanoyl(d31)-2-(9Zoctadecenoyl)-sn-glycero-3-phosphoglycerol (PG(16:0D31/18:1)), 1-hexadecanoyl(d31)-2-(9Z-octadecenoyl)-sn-glycero-3-phospho-(1′-myo-inositol) (PI(16:0D31/18:1)), and 1,1′,2,2′-tetramyristoyl-cardiolipin (sodium salt). Internal standards were added directly to samples prior to MS analysis at a final concentration of 1 μM. The precursor ions of non-oxidized, oxidized and oxidatively truncated PCs in negative-ion mode were detected as formate adducts $[M+HCOO]^-$. Oxidized and oxidatively truncated species demonstrated reproducible retention shifts relative to their non-oxidized precursor lipids and were annotated within defined class-specific retention ranges. Lipid species containing one (+1O) or two (+2O) additional oxygen atoms were assigned as hydroxy- and hydroperoxy-phospholipids, respectively, based on high-resolution MS/MS fragmentation patterns, including precursor ions (detected as $[M–H]^-$ or, for PC species, as formate adducts $[M+HCOO]^-$), diagnostic neutral losses, and characteristic fragment ions consistent with HO- and HOO-containing structures. For low-abundance species with insufficient MS/MS spectra for definitive structural confirmation, annotation was based on accurate mass, retention time, and similarity to structurally validated species. We cannot exclude that a minor fraction of +2O-containing lipid species may represent alternative oxidation isomers (e.g., diols). Low-abundance +3O and +4O-containing phospholipids were also detected, consistent with higher-order oxidation products. Oxidative modifications were localized to the PUFA-containing sn-2 acyl chain based on MS/MS fragmentation. Although oxidation was confirmed by MS/MS, definitive assignment of specific functional groups was not possible for all higher-order species. These likely reflect further oxidation of primary hydroperoxides, yielding complex mixtures of hydroxy-, hydroperoxy-, and epoxy-containing derivatives. Oxidatively truncated phospholipids were identified using a similar multi-criteria approach. Truncated PE, PC, and PS species derived from PUFA-containing sn-2 acyl chains were matched to a curated in-house database based on accurate mass (7 ppm) and class-specific retention time. Structural confirmation was performed by MS/MS, focusing on diagnostic sn-2 fragment ions from an internally generated reference list. Assignments were accepted when predicted truncated acyl fragments, class-specific headgroup ions and other characteristic fragments were detected. Representative annotated MS/MS spectra, including oxidatively truncated phospholipids from each major class, are provided in the Supplementary Figure 6, A-C. Peak areas were used for quantification.

**Lipid quantification and normalization strategy.** Quantification of phospholipid species was performed using a class-based normalization approach. Peak areas were normalized to the corresponding class-specific deuterated internal standards, and calibration curves generated from representative non-oxidized phospholipid standards for each lipid class were used to estimate pmol amounts. Standards used for calibration included:1,2-di-oleoyl-sn-glycero-3-phosphoethanolamine, 1,2-di-oleoyl-sn-glycero-3-phosphocholine, 1,2-di-oleoyl-sn-glycero-3-phosphoserine, 1,2-di-oleoyl-sn-glycero-3-phosphoglycerol, 1,2-di-oleoyl-sn-glycero-3-phosphatidic acid, 1,2-di-oleoyl-sn-glycero-3-phosphoinositol, and 1,1′,2,2′-tetralinoleoyl-cardiolipin. LC-MS data were acquired and analyzed using Xcalibur software. Calculated lipid levels were further normalized to total phospholipid content per sample to account for extraction variability. Because isotope-labeled standards are not available for most truncated and non-truncated oxidized phospholipid species, this strategy provides semi-quantitative estimates rather than absolute molar concentrations. These estimates are appropriate for comparative analysis across experimental conditions and enable robust assessment of relative changes while acknowledging current analytical limitations in redox lipidomics. Aggregated values (e.g., summed HOO-PL or truncated oxidized phospholipids) represent the sum of these semi-quantitative estimates within defined lipid class or oxidation categories and are interpreted in terms of relative differences between groups.

**Statistical Analyses.** All experiments were independently repeated three times, unless otherwise specified in the figure legends. Data are presented as mean ± SD. Statistical analyses were performed using GraphPad Prism 10.2.2 (GraphPad Software Inc.). Comparisons were conducted using an unpaired t-test or one-way ANOVA followed by Tukey’s multiple-comparisons test, as appropriate. $P < 0.05$ was considered statistically significant. Data were graphed using GraphPad Prism 10.2.2 and Origin 2023b. Phospholipids were quantified from full-scan LC–MS spectra using ratiometric comparison to class-specific internal standards and corresponding calibration curves. Principal component analysis (PCA) and orthogonal projections to latent structures–discriminant analysis (OPLS-DA) were performed using SIMCA 18.0 (Sartorius).

**Supplementary Table S1.** Identified oxidatively truncated PE species in HT22 cells

| Exact m/z | Theoretical m/z | ppm error | Formula (as neutral) | Name (CN:DB) | RT, min | MS/MS |
|---|---|---|---|---|---|---|
| **Truncated PE species** | | | | | | |
| 550.3532 | 550.3514 | 3.16 | C27H54O8N1P1 | PE(22:0) | 16.3 | sn-1: m/z 283.2641; sn-2: m/z 87.0455 |
| 562.3152 | 562.3150 | 0.31 | C27H50O9N1P1 | PE(22:2)+1O | 23.6 | sn-1: m/z 281.2489; sn-2: m/z 101.0249 |
| 564.3319 | 564.3307 | 2.13 | C27H52O9N1P1 | PE(22:1)+1O | 23.3 | sn-1: m/z 283.2644; sn-2: m/z 101.0249 |
| 566.3473 | 566.3463 | 1.65 | C27H54O9N1P1 | PE(22:0)+1O | 24.6 | sn-1: m/z 283.2641; sn-2: m/z 103.0402 |
| 576.3312 | 576.3307 | 0.92 | C28H52O9N1P1 | PE(23:2)+1O | 23.6 | sn-1: m/z 283.2639; sn-2: m/z 113.0241 |
| 578.3471 | 578.3463 | 1.27 | C28H54O9N1P1 | PE(23:1)+1O_1 | 20.8 | sn-1: m/z 281.2488; sn-2: m/z 117.0545 |
| 578.3474 | 578.3463 | 1.75 | C28H54O9N1P1 | PE(23:1)+1O_2 | 23.3 | sn-1: m/z 283.2644; sn-2: m/z 115.0392 |
| 578.3829 | 578.3827 | 0.25 | C29H58O8N1P1 | PE(24:0) | 15.0 | sn-1: m/z 283.2639; sn-2: m/z 115.0769 |
| 580.3258 | 580.3256 | 0.27 | C27H52O10N1P1 | PE(22:1)+2O | 23.6 | * |
| 590.3466 | 590.3463 | 0.46 | C29H54O9N1P1 | PE(24:2)+1O | 23.1 | sn-1: m/z 283.2638; sn-2: m/z 127.0406 |
| 592.3636 | 592.3620 | 2.67 | C29H56O9N1P1 | PE(24:1)+1O | 22.8 | sn-1: m/z 283.2634; sn-2: m/z 129.0551 |
| 594.3419 | 594.3413 | 1.04 | C28H54O10N1P1 | PE(23:1)+2O | 24.0 | sn-1: m/z 283.2641; sn-2: m/z 131.0355 |
| 594.3788 | 594.3776 | 1.99 | C29H58O9N1P1 | PE(24:0)+1O | 22.3 | sn-1: m/z 283.2640; sn-2: m/z 131.0719 |
| 602.3482 | 602.3463 | 3.16 | C30H54O9N1P1 | PE(25:3)+1O | 21.1 | sn-1: m/z 255.2331; sn-2: m/z 167.0709 |
| 604.3634 | 604.3620 | 2.30 | C30H56O9N1P1 | PE(25:2)+1O | 24.1 | sn-1: m/z 281.2489; sn-2: m/z 143.0715 |
| 606.3795 | 606.3776 | 3.02 | C30H58O9N1P1 | PE(25:1)+1O | 19.8 | sn-1: m/z 283.2641; sn-2: m/z 143.0698 |
| 616.3635 | 616.3620 | 2.37 | C31H56O9N1P1 | PE(26:3)+1O_1 | 20.0 | sn-1: m/z 255.2335; sn-2: m/z 181.0877 |
| 616.3631 | 616.3620 | 1.87 | C31H56O9N1P1 | PE(26:3)+1O_2 | 20.8 | sn-1: m/z 283.2644; sn-2: m/z 153.0559 |
| 618.3785 | 618.3776 | 1.31 | C31H58O9N1P1 | PE(26:2)+1O | 23.5 | * |
| 620.3586 | 620.3569 | 2.80 | C30H56O10N1P1 | PE(25:2)+2O_1 | 16.5 | sn-1: m/z 283.2640; sn-2: m/z 157.0509 |
| 620.3580 | 620.3569 | 1.82 | C30H56O10N1P1 | PE(25:2)+2O_2 | 25.2 | sn-1: m/z 255.2333; sn-2: m/z 185.0811 |
| 620.3952 | 620.3933 | 3.02 | C31H60O9N1P1 | PE(26:1)+1O | 17.6 | sn-1: m/z 283.2644; sn-2: m/z 157.0877 |
| 622.3759 | 622.3726 | 5.43 | C30H58O10N1P1 | PE(25:1)+2O_1 | 24.6 | sn-1: m/z 281.2488; sn-2: m/z 161.0823 |
| 622.3746 | 622.3726 | 3.32 | C30H58O10N1P1 | PE(25:1)+2O_2 | 25.6 | sn-1: m/z 283.2639; sn-2: m/z 159.0661 |
| 624.3476 | 624.3518 | -6.77 | C29H56O11N1P1 | PE(24:1)+3O | 15.5 | sn-1: m/z 283.2639; sn-2: m/z 161.0456 |
| 630.3798 | 630.3776 | 3.38 | C32H58O9N1P1 | PE(27:3)+1O | 20.2 | sn-1: m/z 283.2639; sn-2: m/z 167.0711 |
| 632.3969 | 632.3933 | 5.68 | C32H60O9N1P1 | PE(27:2)+1O | 17.7 | sn-1: m/z 281.2488; sn-2: m/z 171.1028 |
| 634.3732 | 634.3726 | 0.94 | C31H58O10N1P1 | PE(26:2)+2O | 24.2 | sn-1: m/z 283.2643; sn-2: m/z 171.0666 |
| 634.4122 | 634.4089 | 5.06 | C32H62O9N1P1 | PE(27:1)+1O | 17.1 | sn-1: m/z 283.2646; sn-2: m/z 171.1028 |
| 636.3916 | 636.3882 | 5.26 | C31H60O10N1P1 | PE(26:1)+2O | 24.8 | sn-1: m/z 283.2641; sn-2: m/z 173.0818 |
| 642.3795 | 642.3776 | 2.88 | C33H58O9N1P1 | PE(28:4)+1O | 18.1 | sn-1: m/z 255.2333; sn-2: m/z 207.1028 |
| 644.3951 | 644.3933 | 2.83 | C33H60O9N1P1 | PE(28:3)+1O_1 | 17.8 | sn-1: m/z 255.2336; sn-2: m/z 209.1188 |
| 644.3956 | 644.3933 | 3.64 | C33H60O9N1P1 | PE(28:3)+1O_2 | 19.5 | sn-1: m/z 283.2648; sn-2: m/z 181.0875 |
| 646.3741 | 646.3726 | 2.36 | C32H58O10N1P1 | PE(27:3)+2O | 15.5 | sn-1: m/z 283.2647; sn-2: m/z 183.0669 |
| 648.3909 | 648.3882 | 4.10 | C32H60O10N1P1 | PE(27:2)+2O | 15.3 | sn-1: m/z 283.2641; sn-2: m/z 185.0818 |
| 650.4064 | 650.4039 | 3.89 | C32H62O10N1P1 | PE(27:1)+2O | 23.6 | sn-1: m/z 283.2640; sn-2: m/z 187.0977 |
| 652.3853 | 652.3831 | 3.30 | C31H60O11N1P1 | PE(26:1)+3O | 26.3 | sn-1: m/z 283.2646; sn-2: m/z 189.0766 |
| 656.3967 | 656.3933 | 5.14 | C34H60O9N1P1 | PE(29:4)+1O | 17.3 | sn-1: m/z 283.2641; sn-2: m/z 193.0877 |
| 658.4116 | 658.4089 | 4.04 | C34H62O9N1P1 | PE(29:3)+1O | 16.9 | sn-1: m/z 283.2644; sn-2: m/z 195.1022 |
| 660.3896 | 660.3882 | 2.10 | C33H60O10N1P1 | PE(28:3)+2O | 23.9 | sn-1: m/z 283.2639; sn-2: m/z 197.0820 |
| 664.4220 | 664.4195 | 3.73 | C33H64O10N1P1 | PE(28:1)+2O | 23.9 | * |
| 670.4103 | 670.4089 | 1.99 | C35H62O9N1P1 | PE(30:4)+1O_1 | 16.3 | sn-1: m/z 283.2644; sn-2: m/z 207.1022 |
| 670.4115 | 670.4089 | 3.75 | C35H62O9N1P1 | PE(30:4)+1O_2 | 17.4 | sn-1: m/z 281.2488; sn-2: m/z 209.1188 |
| 672.3903 | 672.3882 | 3.16 | C34H60O10N1P1 | PE(29:4)+2O | 22.5 | sn-1: m/z 255.2333; sn-2: m/z 237.1133 |
| 672.4282 | 672.4246 | 5.43 | C35H64O9N1P1 | PE(30:3)+1O | 17.2 | sn-1: m/z 283.2644; sn-2: m/z 209.1188 |
| 678.4003 | 678.3988 | 2.20 | C33H62O11N1P1 | PE(28:2)+3O | 23.3 | sn-1: m/z 283.2646; sn-2: m/z 215.0931 |

**Extended Supplementary Table S1.** Identified oxidatively truncated PE species in HT22 cells

| Exact m/z | Theoretical m/z | ppm error | Formula (as neutral) | Name (CN:DB) | RT, min | MS/MS |
|---|---|---|---|---|---|---|
| | | **Truncated PE species** | | | | |
| 678.4388 | 678.4352 | 5.38 | C34H66O10N1P1 | PE(29:1)+2O | 18.0 | sn-1: m/z 283.2647; sn-2: m/z 215.1281 |
| 682.4109 | 682.4089 | 2.83 | C36H62O9N1P1 | PE(31:5)+1O | 16.9 | sn-1: m/z 281.2487; sn-2: m/z 221.1188 |
| 684.4280 | 684.4246 | 5.00 | C36H64O9N1P1 | PE(31:4)+1O | 16.6 | sn-1: m/z 283.2646; sn-2: m/z 221.1188 |
| 686.4074 | 686.4039 | 5.21 | C35H62O10N1P1 | PE(30:4)+2O | 16.5 | sn-1: m/z 283.2641; sn-2: m/z 223.0971 |
| 686.4447 | 686.4402 | 6.53 | C36H66O9N1P1 | PE(31:3)+1O | 16.4 | sn-1: m/z 283.2647; sn-2: m/z 223.1344 |
| 688.3845 | 688.3831 | 2.05 | C34H60O11N1P1 | PE(29:4)+3O | 20.9 | sn-1: m/z 283.2640; sn-2: m/z 225.0761 |
| 688.4225 | 688.4195 | 4.28 | C35H64O10N1P1 | PE(30:3)+2O | 20.5 | sn-1: m/z 283.2644; sn-2: m/z 225.1139 |
| 690.4378 | 690.4352 | 3.82 | C35H66O10N1P1 | PE(30:2)+2O | 19.7 | sn-1: m/z 283.2641; sn-2: m/z 227.1279 |
| 694.4324 | 694.4301 | 3.42 | C34H66O11N1P1 | PE(29:1)+3O | 23.9 | sn-1: m/z 283.2640; sn-2: m/z 231.1231 |
| 700.4236 | 700.4195 | 5.89 | C36H64O10N1P1 | PE(31:4)+2O_1 | 18.3 | sn-1: m/z 281.2488; sn-2: m/z 239.1278 |
| 700.4206 | 700.4195 | 1.61 | C36H64O10N1P1 | PE(31:4)+2O_2 | 18.7 | sn-1: m/z 283.2647; sn-2: m/z 237.1133 |
| 702.4022 | 702.3988 | 4.84 | C35H62O11N1P1 | PE(30:4)+3O | 18.4 | * |
| 712.4243 | 712.4195 | 6.79 | C37H64O10N1P1 | PE(32:5)+2O | 19.4 | sn-1: m/z 283.2646; sn-2: m/z 249.1139 |
| 712.4577 | 712.4559 | 2.47 | C38H68O9N1P1 | PE(33:4)+1O | 15.9 | sn-1: m/z 283.2641; sn-2: m/z 249.1499 |
| 714.4379 | 714.4352 | 3.89 | C37H66O10N1P1 | PE(32:4)+2O | 15.9 | * |
| 714.4760 | 714.4715 | 6.30 | C38H70O9N1P1 | PE(33:3)+1O | 15.1 | sn-1: m/z 283.2641; sn-2: m/z 251.1659 |
| 726.4382 | 726.4352 | 4.24 | C38H66O10N1P1 | PE(33:5)+2O | 15.8 | * |
| 728.4186 | 728.4144 | 5.78 | C37H64O11N1P1 | PE(32:5)+3O | 18.3 | sn-1: m/z 283.2647; sn-2: m/z 265.1088 |
| 730.4324 | 730.4301 | 3.22 | C37H66O11N1P1 | PE(32:4)+3O | 16.9 | * |
| 730.4679 | 730.4665 | 2.02 | C38H70O10N1P1 | PE(33:3)+2O | 16.1 | sn-1: m/z 283.2640; sn-2: m/z 267.1600 |

* The precursor ion ($[M-H]^-$) and/or the characteristic headgroup fragments (m/z 140.0092; m/z 196.0335) were detected.
CN - carbon number; DB - double bond number. Repeated annotations represent distinct isomeric or isobaric species, which were resolved by liquid chromatography retention time and confirmed by $MS^2$ fragmentation. In the table, these species are listed in order of increasing retention time (RT) and annotated with suffixes (_1, _2), corresponding to their elution order.

**Supplementary Table S2.** Identified oxidatively truncated PC species in HT22 cells

| Exact m/z | Theoretical m/z | ppm error | Formula (as neutral) | Name (CN:DB) | RT, min | MS/MS |
|---|---|---|---|---|---|---|
| Truncated PC species | | | | | | |
| 610.3762 | 610.3726 | 5.89 | C29H58O10N1P1 | PC(20:0) | 46.14 | * |
| 624.3538 | 624.3518 | 3.18 | C29H56O11N1P1 | PC(20:1)+1O | 59.07 | sn-1: m/z 255.2333; sn-2: m/z 101.0242 |
| 636.3539 | 636.3518 | 3.34 | C30H56O11N1P1 | PC(21:2)+1O | 59.79 | sn-1: m/z 253.2170; sn-2: m/z 115.0400 |
| 636.3900 | 636.3882 | 2.84 | C31H60O10N1P1 | PC(22:1) | 45.59 | [M-$CH_3$-HCOO-H]-; 168.0423 |
| 638.3715 | 638.3675 | 6.26 | C30H58O11N1P1 | PC(21:1)+1O | 56.55 | sn-1: m/z 255.2333; sn-2: m/z 115.0403 |
| 638.4063 | 638.4039 | 3.85 | C31H62O10N1P1 | PC(22:0) | 44.89 | * |
| 650.3691 | 650.3675 | 2.48 | C31H58O11N1P1 | PC(22:2)+1O | 59.29 | * |
| 652.3863 | 652.3831 | 4.86 | C31H60O11N1P1 | PC(22:1)+1O | 53.70 | sn-1: m/z 283.2644; sn-2: m/z 101.0241 |
| 654.4011 | 654.3988 | 3.49 | C31H62O11N1P1 | PC(22:0)+1O | 58.85 | * |
| 664.3877 | 664.3831 | 6.90 | C32H60O11N1P1 | PC(23:2)+1O | 55.49 | sn-1: m/z 281.2480; sn-2: m/z 115.0392 |
| 666.4018 | 666.3988 | 4.55 | C32H62O11N1P1 | PC(23:1)+1O | 55.30 | sn-1: m/z 283.2649; sn-2: m/z 115.0402 |
| 666.4396 | 666.4352 | 6.70 | C33H66O10N1P1 | PC(24:0) | 40.78 | sn-1: m/z 255.2334; sn-2: m/z 143.1077 |
| 678.4379 | 678.4352 | 4.11 | C34H66O10N1P1 | PC(25:1) | 41.02 | * |
| 680.3806 | 680.3780 | 3.82 | C32H60O12N1P1 | PC(23:2)+2O | 52.75 | sn-1: m/z 255.2331; sn-2: m/z 157.0509 |
| 680.4191 | 680.4144 | 6.90 | C33H64O11N1P1 | PC(24:1)+1O | 52.54 | sn-1: m/z 255.2329; sn-2: m/z 157.0871 |
| 690.4023 | 690.3988 | 5.18 | C34H62O11N1P1 | PC(25:3)+1O | 51.68 | sn-1: m/z 255.2327; sn-2: m/z 167.0711 |
| 692.4174 | 692.4144 | 4.28 | C34H64O11N1P1 | PC(25:2)+1O_1 | 49.79 | sn-1: m/z 255.2332; sn-2: m/z 169.0871 |
| 692.4180 | 692.4144 | 5.19 | C34H64O11N1P1 | PC(25:2)+1O_2 | 50.98 | sn-1: m/z 281.2483; sn-2: m/z 143.0712 |
| 692.4166 | 692.4144 | 3.08 | C34H64O11N1P1 | PC(25:2)+1O_3 | 55.45 | * |
| 692.4549 | 692.4508 | 5.93 | C35H68O10N1P1 | PC(26:1) | 40.12 | sn-1: m/z 255.2326; sn-2: m/z 169.1233 |
| 694.4340 | 694.4301 | 5.71 | C34H66O11N1P1 | PC(25:1)+1O_1 | 50.09 | sn-1: m/z 255.2329; sn-2: m/z 171.1016 |
| 694.4344 | 694.4301 | 6.24 | C34H66O11N1P1 | PC(25:1)+1O_2 | 51.99 | sn-1: m/z 283.2639; sn-2: m/z 143.0715 |
| 696.4131 | 696.4093 | 5.37 | C33H64O12N1P1 | PC(24:1)+2O | 46.97 | sn-1: m/z 255.2322; sn-2: m/z 173.0816 |
| 704.4187 | 704.4144 | 6.11 | C35H64O11N1P1 | PC(26:3)+1O | 51.50 | sn-1: m/z 255.2329; sn-2: m/z 181.0879 |
| 706.4342 | 706.4301 | 5.89 | C35H66O11N1P1 | PC(26:2)+1O | 47.93 | sn-1: m/z 255.2326; sn-2: m/z 183.1028 |
| 708.4134 | 708.4093 | 5.74 | C34H64O12N1P1 | PC(25:2)+2O | 50.46 | sn-1: m/z 283.2646; sn-2: m/z 157.0509 |
| 708.4491 | 708.4457 | 4.80 | C35H68O11N1P1 | PC(26:1)+1O | 47.63 | sn-1: m/z 255.2333; sn-2: m/z 185.1188 |
| 712.4444 | 712.4406 | 5.25 | C34H68O12N1P1 | PC(25:0)+2O | 56.00 | sn-1: m/z 255.2334; sn-2: m/z 189.1134 |
| 716.4179 | 716.4144 | 4.85 | C36H64O11N1P1 | PC(27:4)+1O_1 | 48.25 | sn-1: m/z 255.2335; sn-2: m/z 193.0871 |
| 716.4187 | 716.4144 | 6.02 | C36H64O11N1P1 | PC(27:4)+1O_2 | 51.88 | sn-1: m/z 281.2489; sn-2: m/z 167.0712 |
| 718.4350 | 718.4301 | 6.81 | C36H66O11N1P1 | PC(27:3)+1O_1 | 47.10 | sn-1: m/z 255.2333; sn-2: m/z 195.1022 |
| 718.4342 | 718.4301 | 5.70 | C36H66O11N1P1 | PC(27:3)+1O_2 | 50.22 | sn-1: m/z 283.2646; sn-2: m/z 167.0710 |
| 720.4114 | 720.4093 | 2.93 | C35H64O12N1P1 | PC(26:3)+2O | 59.20 | sn-1: m/z 255.2336; sn-2: m/z 197.0818 |
| 720.4490 | 720.4457 | 4.53 | C36H68O11N1P1 | PC(27:2)+1O | 49.03 | sn-1: m/z 283.2645; sn-2: m/z 169.0877 |
| 730.4335 | 730.4301 | 4.74 | C37H66O11N1P1 | PC(28:4)+1O | 50.39 | sn-1: m/z 281.2487; sn-2: m/z 181.0871 |
| 730.4693 | 730.4665 | 3.87 | C38H70O10N1P1 | PC(29:3) | 44.64 | sn-1: m/z 255.2321; sn-2: m/z 207.1399 |
| 732.4125 | 732.4093 | 4.29 | C36H64O12N1P1 | PC(27:4)+2O | 56.38 | sn-1: m/z 255.2335; sn-2: m/z 209.0817 |
| 732.4506 | 732.4457 | 6.72 | C37H68O11N1P1 | PC(28:3)+1O_1 | 47.12 | sn-1: m/z 255.2337; sn-2: m/z 209.1188 |
| 732.4505 | 732.4457 | 6.53 | C37H68O11N1P1 | PC(28:3)+1O_2 | 49.34 | sn-1: m/z 283.2648; sn-2: m/z 181.0879 |
| 742.4349 | 742.4301 | 6.46 | C38H66O11N1P1 | PC(29:5)+1O | 47.15 | sn-1: m/z 281.2480; sn-2: m/z 193.0869 |
| 744.4493 | 744.4457 | 4.87 | C38H68O11N1P1 | PC(29:4)+1O | 46.37 | sn-1: m/z 283.2638; sn-2: m/z 193.0875 |
| 756.4482 | 756.4457 | 3.31 | C39H68O11N1P1 | PC(30:5)+1O | 44.10 | * |
| 758.4631 | 758.4614 | 2.23 | C39H70O11N1P1 | PC(30:4)+1O | 46.11 | sn-1: m/z 281.2488; sn-2: m/z 193.0871 |
| 770.4651 | 770.4614 | 4.83 | C40H70O11N1P1 | PC(31:5)+1O | 45.11 | sn-1: m/z 281.2486; sn-2: m/z 221.1188 |
| 772.4807 | 772.4770 | 4.70 | C40H72O11N1P1 | PC(31:4)+1O | 44.42 | sn-1: m/z 283.2639; sn-2: m/z 221.1179 |

* The precursor ion ($[M+HCOO]^-$ and/or $[M-CH_3-H]^-$) and/or the characteristic phosphocholine headgroup fragment (m/z 168.0431) were detected.
CN - carbon number; DB - double bond number. Repeated annotations represent distinct isomeric or isobaric species, which were resolved by liquid chromatography retention time and confirmed by MS² fragmentation. In the table, these species are listed in order of increasing retention time (RT) and annotated with suffixes (_1, _2, _3), corresponding to their elution order.

**Supplementary Table S3.** Identified oxidatively truncated PS species in HT22 cells

| Exact m/z | Theoretical m/z | ppm error | Formula (as neutral) | Name (CN:DB) | RT, min | MS/MS |
|---|---|---|---|---|---|---|
| **Truncated PS species** | | | | | | |
| 594.3436 | 594.3413 | 3.87 | C28H54O10N1P1 | PS(22:0) | 33.10 | sn-1: m/z 283.2631; sn-2: m/z 87.0450 |
| 634.3734 | 634.3726 | 1.26 | C31H58O10N1P1 | PS(25:1) | 32.11 | * |
| 636.3540 | 636.3518 | 3.46 | C30H56O11N1P1 | PS(24:1)+1O | 31.13 | sn-1: m/z 283.2640; sn-2: m/z 129.0553 |
| 638.3711 | 638.3675 | 5.64 | C30H58O11N1P1 | PS(24:0)+1O | 30.64 | sn-1: m/z 283.2640; sn-2: m/z 131.0711 |
| 650.3714 | 650.3675 | 6.00 | C31H58O11N1P1 | PS(25:1)+1O | 30.55 | sn-1: m/z 283.2636; sn-2: m/z 143.0699 |
| 668.3810 | 668.3780 | 4.49 | C31H60O12N1P1 | PS(25:0)+2O | 31.13 | sn-1: m/z 283.2639; sn-2: m/z 161.0822 |
| 678.4035 | 678.3988 | 6.93 | C33H62O11N1P1 | PS(27:1)+1O | 30.21 | sn-1: m/z 283.2642; sn-2: m/z 171.1023 |
| 688.3842 | 688.3831 | 1.60 | C34H60O11N1P1 | PS(28:3)+1O | 30.25 | * |
| 716.4146 | 716.4144 | 0.28 | C36H64O11N1P1 | PS(30:3)+1O | 30.01 | sn-1: m/z 283.2634; sn-2: m/z 209.1174 |
| 732.4100 | 732.4093 | 0.96 | C36H64O12N1P1 | PS(30:3)+2O | 30.74 | sn-1: m/z 283.2639; sn-2: m/z 225.1130 |
| 738.4235 | 738.4199 | 4.88 | C35H66O13N1P1 | PS(29:1)+3O | 31.35 | sn-1: m/z 283.2637; sn-2: m/z 231.1236 |

* The precursor ion ($[M\text{-}H]^-$) and/or the precursor ion with loss of the serine headgroup ($[M\text{-}C_3H_7NO_3\text{-}H]^-$) were detected.
CN - carbon number; DB - double bond number.

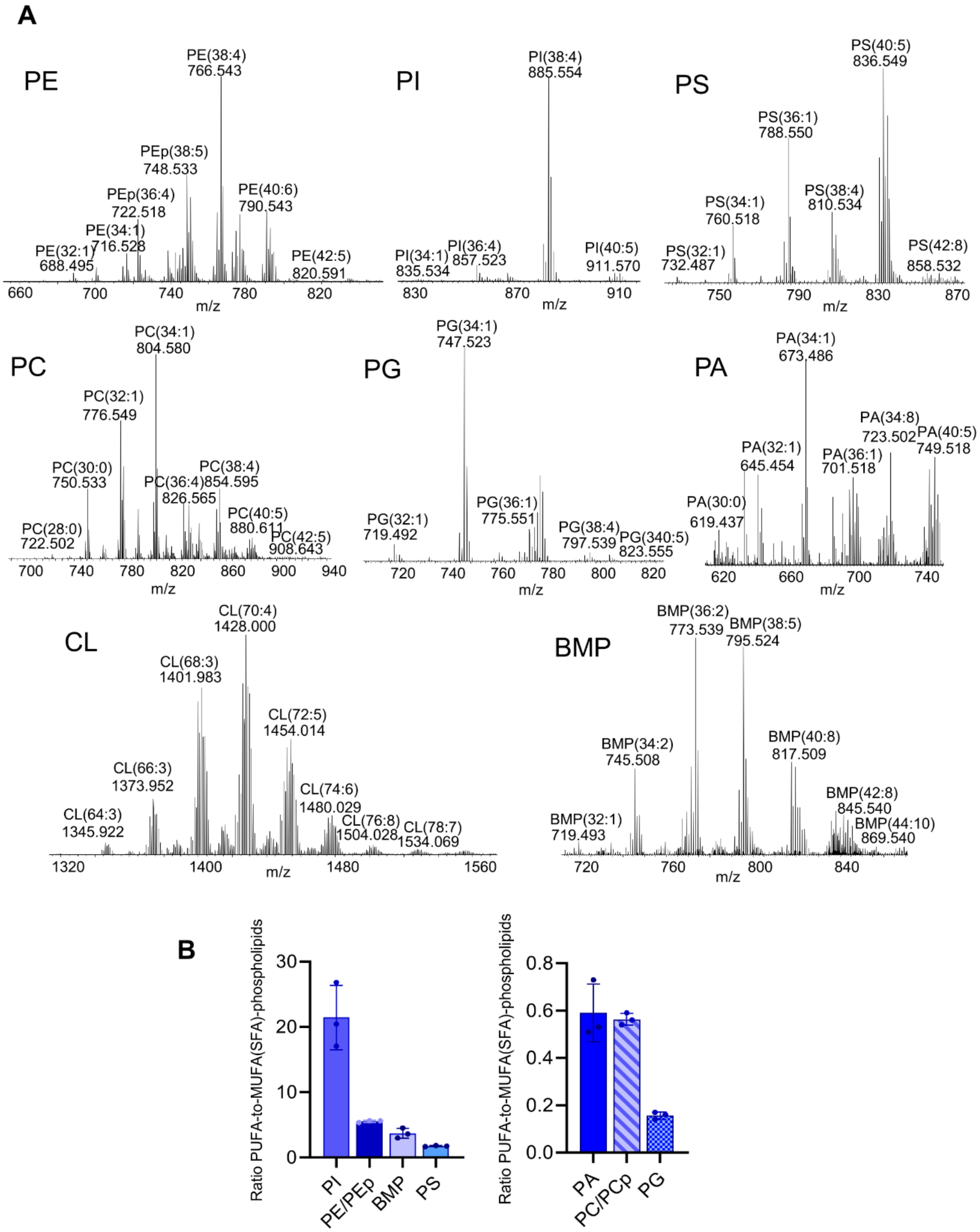


**Supplementary Figure S1. LC-MS analysis of the phospholipidome in HT22 cells. (A)** Representative mass spectra of major phospholipids from HT22 cells obtained using normal-phase chromatography, corresponding to MS1 scans acquired at the retention times where specific lipid classes are expected to elute. Lipid species are annotated as PL(CN:DB), where CN is the total number of carbon atoms and DB is the total number of double bonds in the combined sn-1 and sn-2 acyl chains. PLs, phospholipids; CL, cardiolipin; PC, phosphatidylcholine; PE, phosphatidylethanolamine; PS, phosphatidylserine; PI, phosphatidylinositol; PA, phosphatidic acid; PG, phosphatidylglycerol; BMP, bis-monoacylglycero-phosphate; OxTr PLs, oxidatively truncated phospholipids. **(B)** Comparison of PUFA- versus MUFA/SFA-containing phospholipids. Data are mean±SD, n = 3/group. Note: two plots are shown to accommodate different abundance ranges.

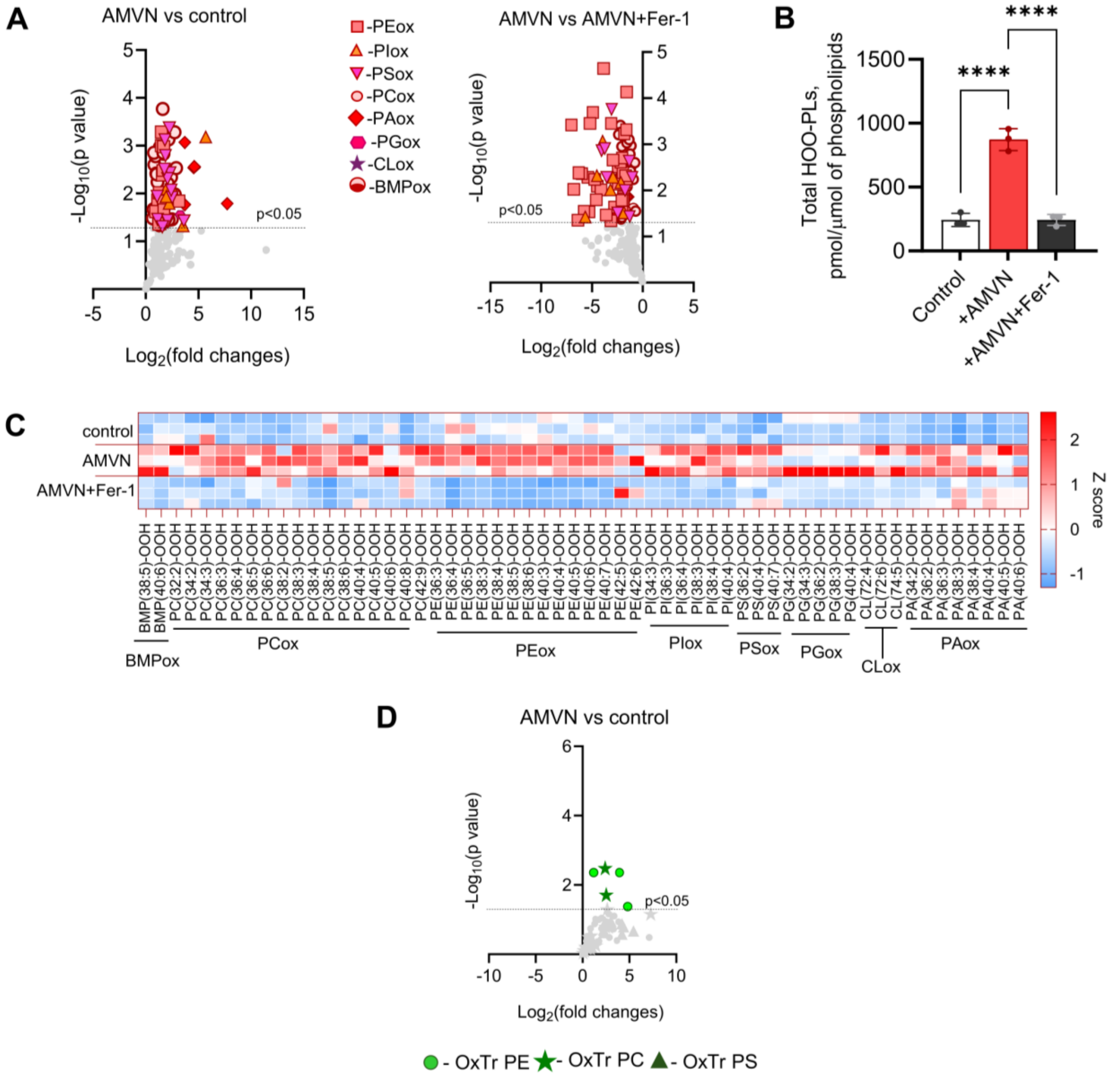


**Supplementary Figure S2. Ferroptosis-independent lipid peroxidation induced by AMVN in HT22 cells. (A)** Volcano plots comparing AMVN vs. control (left) and AMVN vs. AMVN+Fer-1 (right) for all oxidized PL species (containing 1-4 oxygens), highlighting significantly altered species. **(B)** Quantification of total HOO-PLs (pmol/μmol of phospholipids) in HT22 cells under control, AMVN, and AMVN+Fer-1 treatments. Data are shown as mean±SD, n = 3/group, Ordinary one-way ANOVA test, ****p < 0.0001. **(C)** Heat map showing the relative abundances of HOO-PLs in HT22 cells under control, AMVN, and AMVN+Fer-1 treatments. Values are expressed as pmol/μmol of phospholipids (n = 3/group) and displayed as z-score–scaled heat maps, color-coded from blue (low) to red (high). **(D)** Volcano plot showing significantly altered oxidatively truncated phospholipid species (truncated PE, PC, and PS) in AMVN-treated vs. control cells.

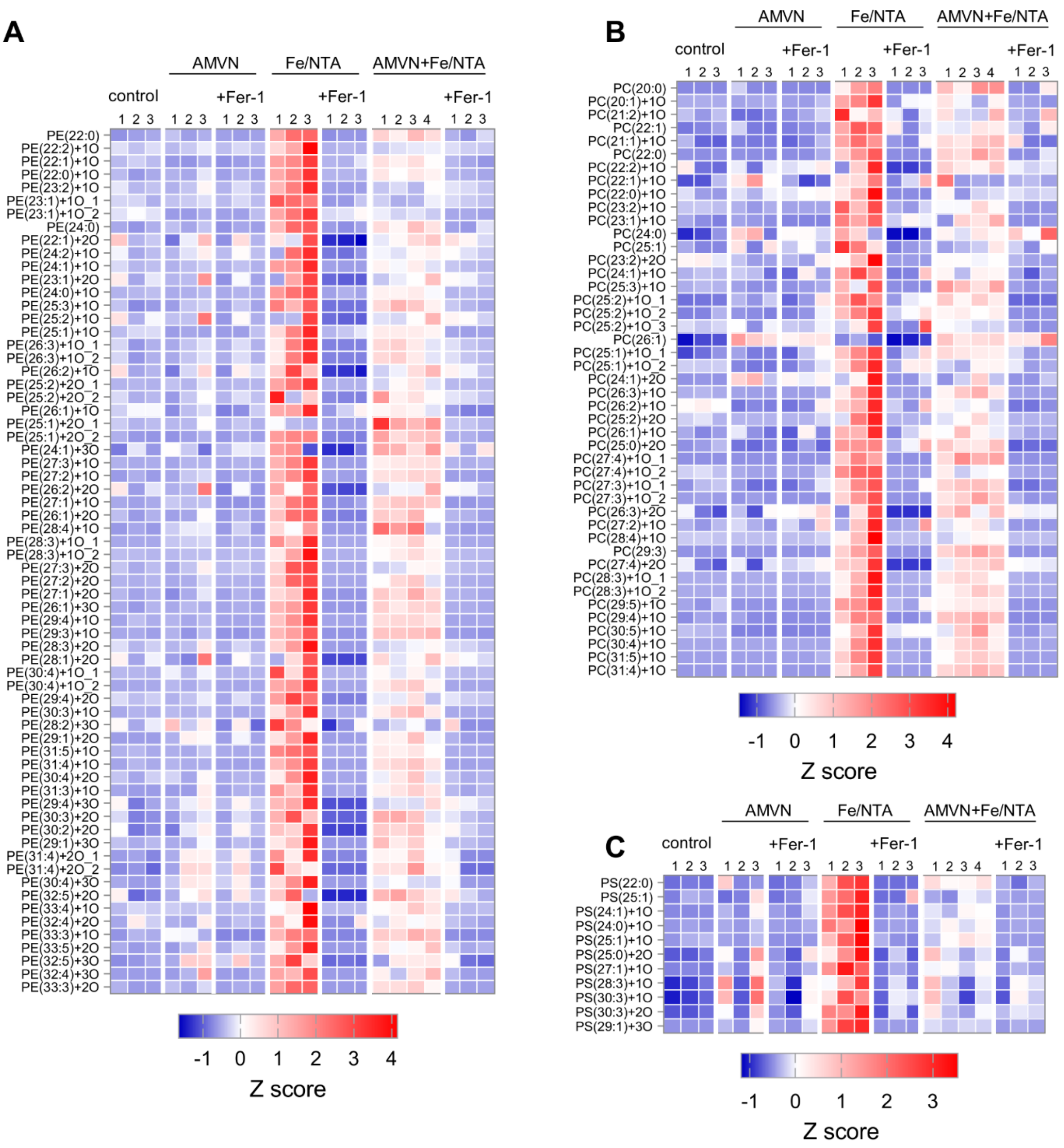


**Supplementary Figure S3. Oxidatively truncated phospholipid profiles across PE, PC, and PS classes in HT22 cells.** Heat maps showing the relative abundance of oxidatively truncated phospholipids species in HT22 cells under control, AMVN±Fer-1, Fe/NTA±Fer-1, and AMVN±Fer-1 treatments. Values are expressed as pmol/μmol of phospholipids (n = 3-4/group) and displayed as z-score–scaled heat maps, color-coded from blue (low) to red (high). **(A)** Phosphatidylethanolamine (PE), **(B)** Phosphatidylcholine (PC), and **(C)** Phosphatidylserine (PS) classes. Truncated lipid species are annotated as PL(CN:DB), where CN is the total number of carbon atoms and DB is the total number of double bonds in the combined sn-1 and sn-2 acyl chains, with or without additional oxygen(s) (+1O, +2O, or +3O). Repeated annotations represent distinct isomeric/isobaric species, resolved by LC retention time and confirmed by MS², listed by increasing retention time and annotated with suffixes (_1, _2, _3) for elution order.

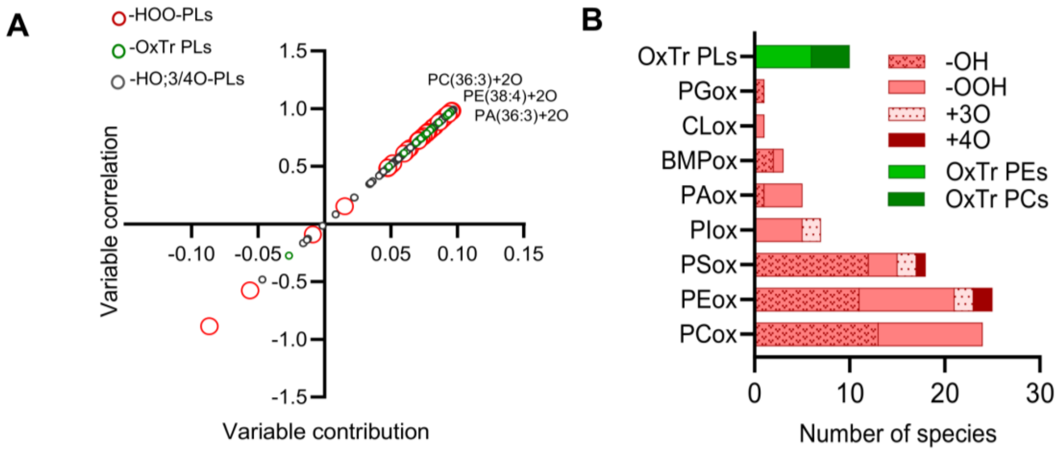


**C**

| Species | Fold change | p value | Rank | VIPpred score |
|---|---|---|---|---|
| PC(36:6)-OH | 3. 1 | 0.0002 | 1 | 1.3003 |
| PS(38:4)-OH | 4.7 | 0.0004 | 2 | 1.2892 |
| PE(40:6)-OH | 2.8 | 0.0005 | 3 | 1.2881 |
| PC(36:3)-OOH | 6.7 | 0.0005 | 4 | 1.2868 |
| PI(40:4)+3O | 51.3 | 0.0007 | 5 | 1.2860 |
| PC(36:2)-OH | 4.5 | 0.0007 | 6 | 1.2837 |
| PE(38:4)-OOH | 2.8 | 0.0010 | 7 | 1.2831 |
| PS(36:2)-OH | 3.5 | 0.0008 | 8 | 1.2812 |
| PA(36:3)-OOH | 13.1 | 0.0009 | 9 | 1.2806 |
| PE(40:4)-OH | 3.2 | 0.0011 | 10 | 1.2805 |

**Supplementary Figure S4. Group-separating lipid species in AMVN-treated HT22 cells. (A)** OPLS-DA-derived S-plot for the AMVN treatment. **(B)** Distribution of the 94 group-separating species across oxidized and OxTr phospholipid classes, based on VIP scores (VIPpred > 1). **(C)** Top 10 ranked group-separating species for the AMVN treatment based on VIPpred score, fold change relative to control, and $p < 0.05$.

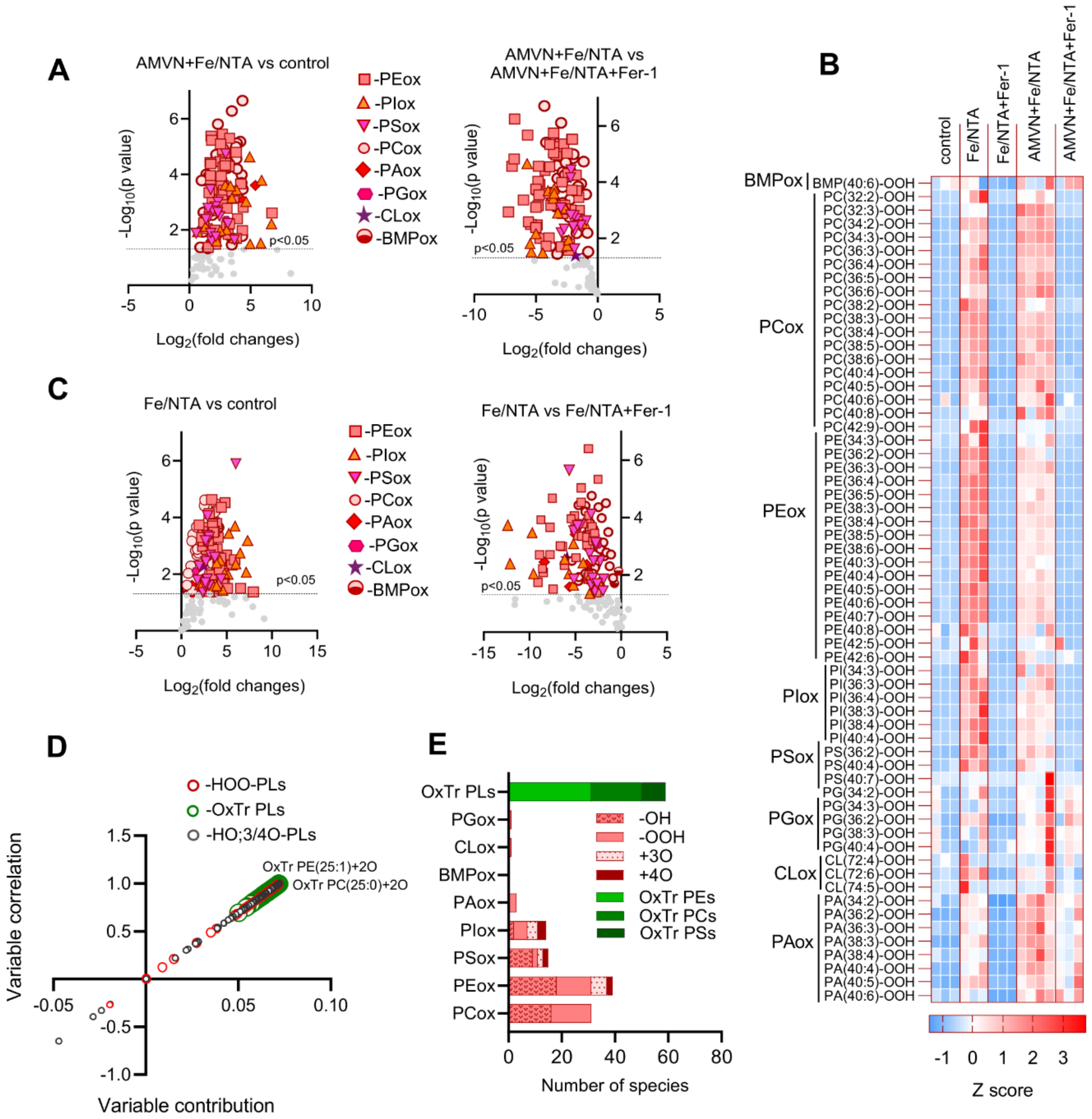


**F**

| Species | Fold change | p value | Rank | VIPpred score |
|---|---|---|---|---|
| PS(40:3)+4O | 64.5 | <0.0001 | 1 | 1.150 |
| PE(36:3)-OH | 10.2 | <0.0001 | 2 | 1.146 |
| PC(36:5)-OH | 5.9 | <0.0001 | 3 | 1.146 |
| PE(36:5)+3O | 27.8 | <0.0001 | 4 | 1.146 |
| PC(34:2)-OH | 13.6 | <0.0001 | 5 | 1.145 |
| PC(36:3)-OH | 5.7 | <0.0001 | 6 | 1.145 |
| OxTr PC(25:0)+2O | 5.8 | <0.0001 | 7 | 1.145 |
| PE(38:7)-OH | 21.4 | <0.0001 | 8 | 1.145 |
| OxTr PE(25:1)+2O | 8.9 | 0.0001 | 9 | 1.144 |
| PC(34:2)-OOH | 13.9 | 0.0001 | 10 | 1.144 |

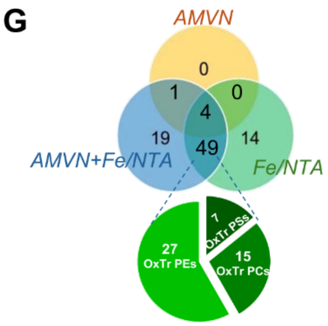


**Supplementary Figure S5. Expanded characterization and supporting multivariate analyses of oxidized phospholipids in AMVN+Fe/NTA- and Fe/NTA-treated HT22 cells. (A)** Volcano plots comparing AMVN+Fe/NTA vs. control (left) and AMVN+Fe/NTA +Fer-1 vs. AMVN+Fe/NTA (right) for all oxidized phospholipid classes, highlighting significantly altered species. **(B)** Heat map showing HOO-PL abundances in HT22 cells under control, AMVN+Fe/NTA± Fer-1, and Fe/NTA± Fer-1 treatments. Values are expressed as pmol/μmol of phospholipids (n = 3-4/group) and displayed as z-score–scaled heat maps, color-coded from blue (low) to red (high). **(C)** Volcano plots comparing Fe/NTA vs. control (left) and Fe/NTA+Fer-1 vs. Fe/NTA (right) for all oxidized phospholipid classes, highlighting significantly altered species. **(D)** OPLS-DA-derived S-plot for the Fe/NTA treatment. **(E)** Distribution of the 162 group-separating species across oxidized and OxTr phospholipid species, based on VIP scores (VIPpred > 1). **(F)** Top 10 ranked group-separating species for the Fe/NTA treatment based on VIP score, fold change relative to control, and $p < 0.05$. Note: OxTr-PE(25:1)+2O refers to the species eluting at RT = 25.6 min; see Supplementary Table S1. **(G)** Venn diagram comparing significantly altered oxidatively truncated phospholipid species across AMVN, Fe/NTA, and AMVN+Fe/NTA treatments. The insert pie chart shows the distribution of truncated PE, PC, and PS species among OxTr phospholipids common to Fe/NTA and AMVN+Fe/NTA, with OxTr PEs representing the majority (55.1%).

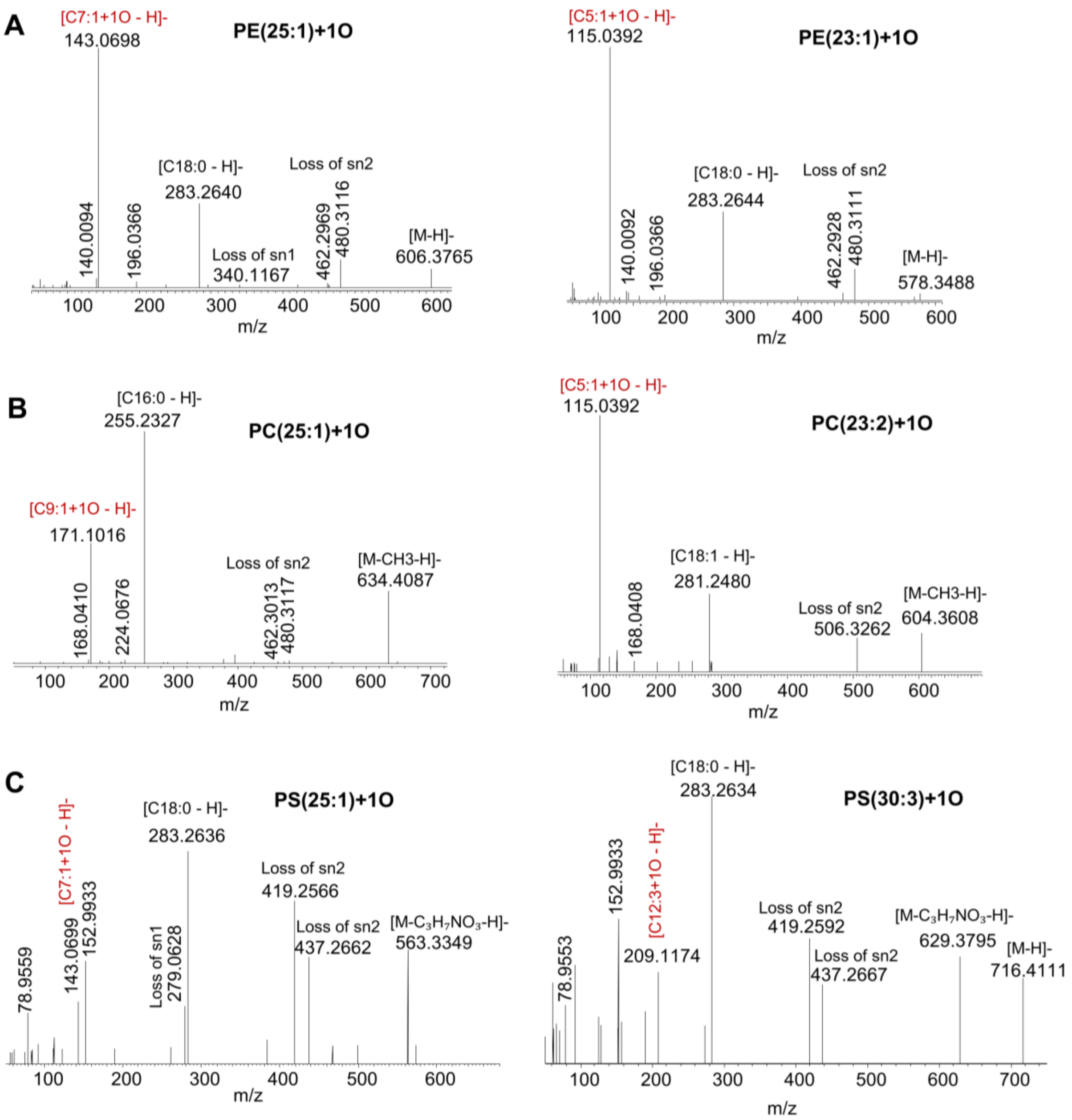


**Supplementary Figure S6. Representative MS/MS spectra showing PE, PC, and PS truncated phospholipid species. (A)** *Left*. MS/MS spectrum of PE(18:0/7:1)+1O, [M-H]- at m/z 606.3765. sn-1 localization of 18:0 is supported by the acyl fragment at m/z 283.2640 and lyso-PE ions at m/z 462.2969 and m/z 480.3116 (neutral loss of sn-2 fatty acyl chain as free fatty acid (FFA) and ketene, respectively). The fragment ion at m/z 143.0668 is annotated as a putative C7-truncated sn-2 fatty acyl fragment (C7:1+1O) and is further supported by the lyso-PE ion at m/z 340.1167 (the loss of the sn-1 fatty acyl chain). *Right*. MS/MS spectrum of PE(18:0/5:1)+1O, [M-H]- at m/z 578.3488. sn-1 localization of 18:0 is supported by the acyl fragment at m/z 283.2644 and lyso-PE ions at m/z 462.2928 and m/z 480.3111 (neutral loss of the sn-2 fatty acyl chain as FFA and ketene, respectively). The fragment ion at m/z 115.0372 is annotated as a putative C5-truncated sn-2 fatty acyl fragment (C5:1+1O). **(B)** *Left*. MS/MS spectrum of PC(16:0/9:1)+1O, [M-CH3-H]- at m/z 634.4087. sn-1 localization to 16:0 is supported by the acyl fragment at m/z 255.2327 and lyso-PC ions at m/z 462.3013 (neutral loss of the sn-2 fatty acyl chain as FFA, with loss of CH3 and formate from the precursor ion) and m/z 480.3117 (loss of the sn-2 fatty acyl chain as ketene, CH3 and formate from the precursor ion). The fragment ion at m/z 171.1006 is annotated as a putative C9-truncated sn-2 fatty acyl fragment (C9:1+1O). *Right*. MS/MS spectrum of PC(18:1/5:1)+1O, [M-CH3-H]- at m/z 604.3608. sn-1 localization of 18:1 is supported by the acyl fragment at m/z 281.2480 and lyso-PC ion at m/z 506.3262 (loss of the sn-2 fatty acyl chain as ketene, CH3 and formate from the precursor ion). The fragment ion at m/z 115.0372 is annotated as a putative C5-

truncated sn-2 fatty acyl fragment (C5:1+1O). **(C)** *Left*. MS/MS spectrum of PS (18:0/7:1)+1O, [M-$C_3H_7NO_3$-H]- (loss of serine group) at m/z 563.3349. sn-1 localization of 18:0 is supported by the acyl fragment at m/z 283.2636 and lyso-PS ions at m/z 419.2566 and m/z 437.2662 (neutral loss of the sn-2 fatty acyl chain as FFA and ketene with serine from the precursor ion, respectively). The fragment ion at m/z 143.0690 is annotated as a putative C7-truncated sn-2 fatty acyl fragment (C7:1+1O) and is further supported by the lyso-PS ion at m/z 279.0628 (the loss of the sn-1 fatty acyl chain). *Right*. MS/MS spectrum of PS(18:0/12:2)+1O, [M-H]- at m/z 716.4111. sn-1 localization of 18:0 is supported by the acyl fragment at m/z 283.2634 and lyso-PS ions at m/z 419.2592 and m/z 437.2667 (neutral loss of the sn-2 fatty acyl chain as FFA and ketene with serine from the precursor ion, respectively). The fragment ion at m/z 209.1164 is annotated as a putative C12-truncated sn-2 fatty acyl fragment (C12:3+1O).

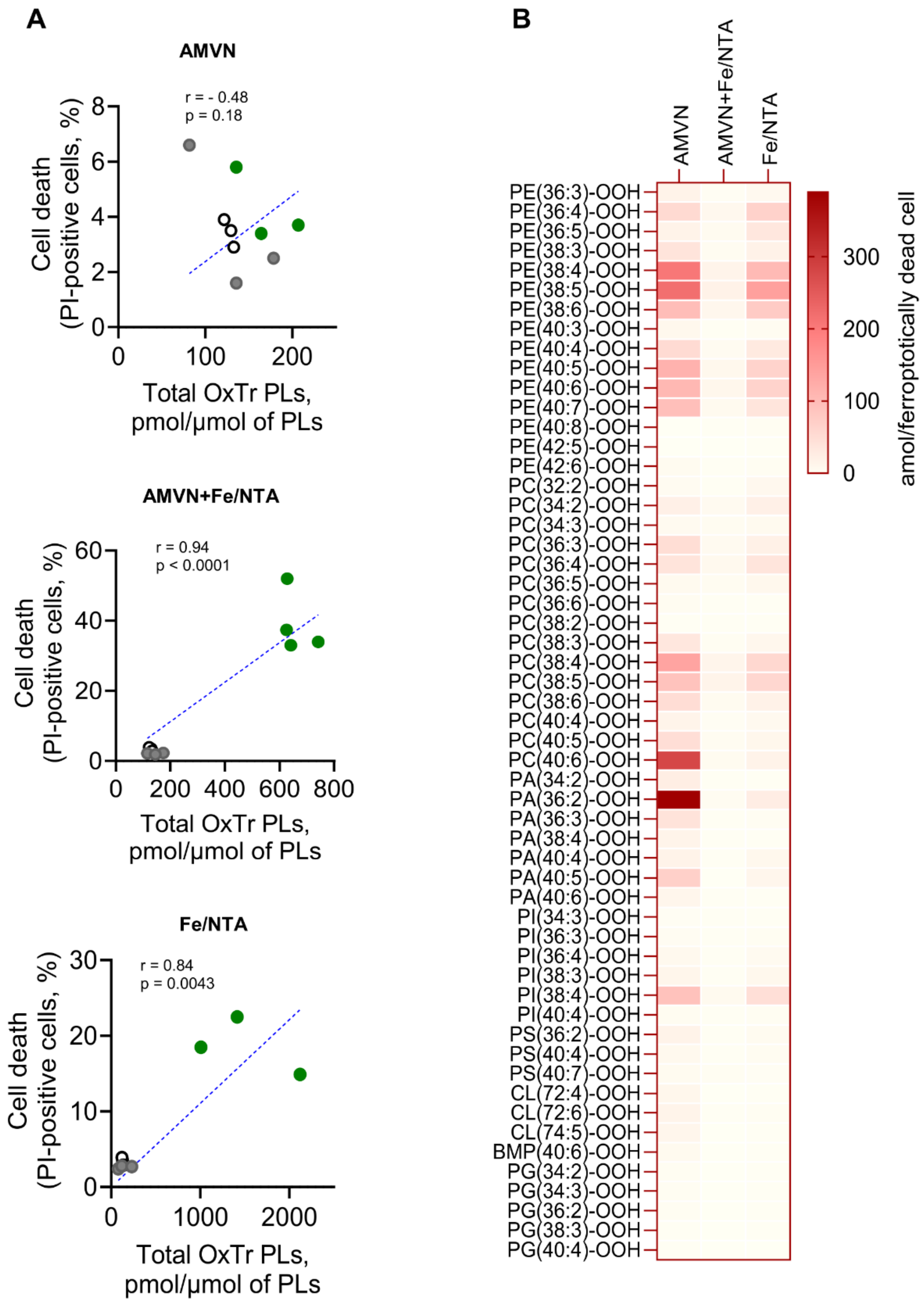


**Supplementary Figure S7. Correlation of oxidatively truncated phospholipids with cell death and hydroperoxy-phospholipid burden per dying cell under different conditions. (A)** Pearson correlation plots of total OxTr PLs with cell death for (upper) control, AMVN ± Fer-1; (middle) control, AMVN+Fe/NTA ± Fer-1; and (lower) control, Fe/NTA ± Fer-1 treatments in HT22 cells. Each dot represents an independent replicate. **(B)** Heat map of individual HOO-PL species across different lipid classes for three experimental conditions (AMVN, Fe/NTA, AMVN+Fe/NTA), showing HOO-PL levels after subtraction of Fer-1-insensitive background, which were normalized to the number of Fer-1 sensitive dead cells to estimate the hydroperoxide burden. Data are presented as mean (n = 3-4/group) in amol per ferroptosis-attributable cell death.

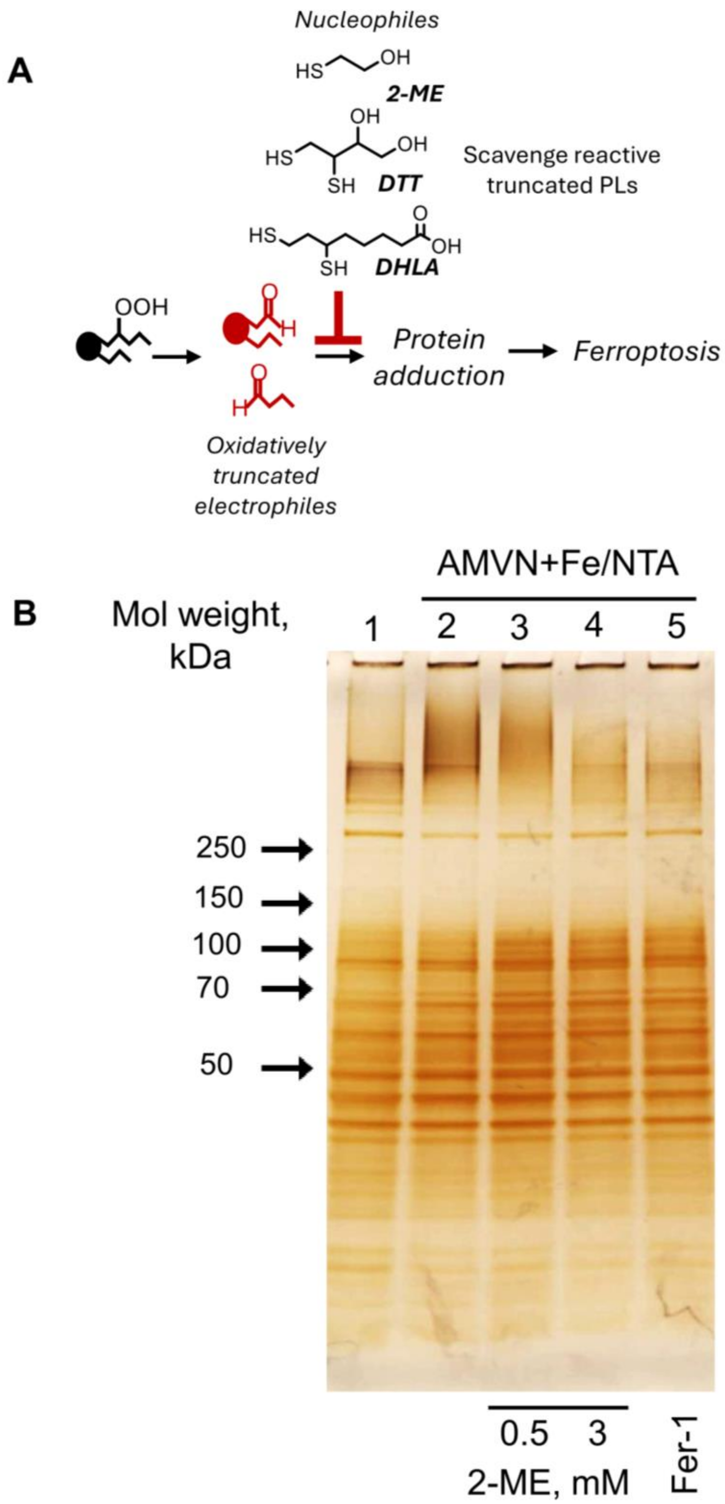


**Supplementary Figure S8. Protective effects of nucleophiles on truncated lipid-protein adduct formation in HT22 cells. (A)** Schematic representation illustrating how nucleophilic molecules, 2-mercaptoethanol (ME), dithiothreitol (DTT), and dihydrolipoic acid (DHLA), can react with truncated electrophilic lipid species to protect from lipid-protein adduct formation and ferroptotic cell death. **(B)** Cell lysates from control (1), AMVN+Fe/NTA (2), AMVN+Fe/NTA with 0.5 mM 2-ME (3), AMVN+Fe/NTA with 3 mM 2-ME (4), and AMVN+Fe/NTA with Fer-1 (5) were analyzed by PAGE followed by silver staining. Shown is a representative gel from n = 3-7 independent experiments.
